# Modulation of the promoter activation rate dictates the transcriptional response to graded BMP signaling levels in the *Drosophila* embryo

**DOI:** 10.1101/837179

**Authors:** Caroline Hoppe, Jonathan R. Bowles, Thomas G. Minchington, Catherine Sutcliffe, Priyanka Upadhyai, Magnus Rattray, Hilary L. Ashe

## Abstract

Morphogen gradients specify cell fates during development, with a classic example being the BMP gradient’s conserved role in embryonic dorsal-ventral axis patterning. Here we elucidate how the BMP gradient is interpreted in the *Drosophila* embryo, by combining live imaging with computational modelling to infer transcriptional burst parameters at single cell resolution. By comparing burst kinetics in cells receiving different levels of BMP signaling, we show that BMP signaling controls burst frequency by regulating the promoter activation rate. We provide evidence that the promoter activation rate is influenced by both enhancer and promoter sequences, whereas Pol II loading rate is primarily modulated by the enhancer. Consistent with BMP-dependent regulation of burst frequency, the numbers of BMP target gene transcripts per cell are graded across their expression domains. We suggest that graded mRNA output is a general feature of morphogen gradient interpretation and discuss how this can impact on cell fate decisions.

## INTRODUCTION

A gradient of Bone Morphogenetic Protein (BMP) signaling patterns ectodermal cell fates along the dorsal-ventral axis of vertebrate and invertebrate embryos (Bier and De Robertis, 2015; Hamaratoglu et al., 2014). In *Drosophila*, visualization of Decapentaplegic (Dpp), the major BMP signaling molecule, reveals a shallow graded distribution in early embryos that subsequently refines to a peak of Dpp at the dorsal midline (Shimmi et al., 2005; Wang and Ferguson, 2005). BMP-receptor activation leads to phosphorylation of the Mothers against dpp (Mad) transcription factor, which associates with Medea (Med) to activate or repress target gene transcription (Hamaratoglu et al., 2014). A stripe of phosphorylated Mad (pMad) and Med centered at the dorsal midline has been visualized in the early *Drosophila* embryo (Dorfman and Shilo, 2001; Rushlow et al., 2001; Sutherland et al., 2003), similar to that observed for Dpp (Shimmi et al., 2005; Wang and Ferguson, 2005), although lower pMad levels are also detectable in a few adjacent dorsal-lateral cells (Rushlow et al., 2001). The BMP/pMad gradient activates different thresholds of gene activity, including the peak target gene *hindsight* (*hnt*) and the intermediate target *u-shaped* (*ush*) (Ashe et al., 2000).

New insights into transcriptional activation have been obtained by studying this process in single cells using quantitative and live imaging approaches, including single molecule FISH (smFISH) and the MS2/MCP system (Pichon et al., 2018). The latter allowed the first direct visualization of pulses or bursts of transcriptional activity (Chubb et al., 2006; Golding et al., 2005). Enhancers have been shown to regulate the frequency of transcriptional bursts, with strong enhancers generating more bursts than weaker enhancers (Fukaya et al., 2016; Larson et al., 2013; Larsson et al., 2019; Senecal et al., 2014). In addition, the detection of simultaneous bursts of transcription of two linked reporters by a single enhancer argues against the classic enhancer-promoter looping model (Fukaya et al., 2016).

Recently, Notch target genes in *Drosophila* and *C. elegans* have been shown to undergo transcriptional bursting, with Notch controlling burst size through effects on duration (Falo-Sanjuan et al., 2019; Lee et al., 2019). However, it is unclear whether this is a general mechanism by which signals control transcriptional bursting. Therefore, to provide insight into BMP gradient interpretation at single cell resolution, we have used live imaging and quantitative analysis to determine the kinetics of endogenous BMP target gene transcription in the *Drosophila* embryo. These data reveal that BMP signaling modulates the promoter activation rate and therefore burst frequency. The different burst frequencies of BMP target genes, depending on cellular position, result in a gradient of mRNA numbers per cell. Overall these data reveal how a signaling gradient is decoded with different transcriptional kinetics to impart positional information on cells.

## RESULTS

### Temporal dynamics of transcriptional activation in response to the BMP gradient

We used the MS2 system (Garcia et al., 2013; Lucas et al., 2013) to visualize the temporal dynamics of BMP-responsive transcription during early embryogenesis. CRISPR genome engineering was used to introduce 24 copies of the MS2 stem loops into the endogenous 5’ UTR of the *hnt* and *ush* genes, which respond to peak and intermediate BMP signaling levels, respectively (Ashe et al., 2000) (Fig. 1A). Conventional in situ hybridization showed *ush* and *hnt* expression patterns equivalent to those observed in wildtype (wt) embryos (Fig. S1A), indicating that insertion of the loops does not affect the expression patterns. To visualize transcription dynamics, females maternally expressing one copy of MCP-GFP, which binds to the MS2 loops, and Histone-RFP were crossed to males carrying the *ush* or *hnt* gene with MS2 stem loops, so the resulting embryos have a single allele carrying the MS2 sequence. Confocal imaging of these embryos allows the bright fluorescent signal associated with the nascent transcription site to be recorded for each expressing nucleus (Fig. 1B).

**Figure 1:**
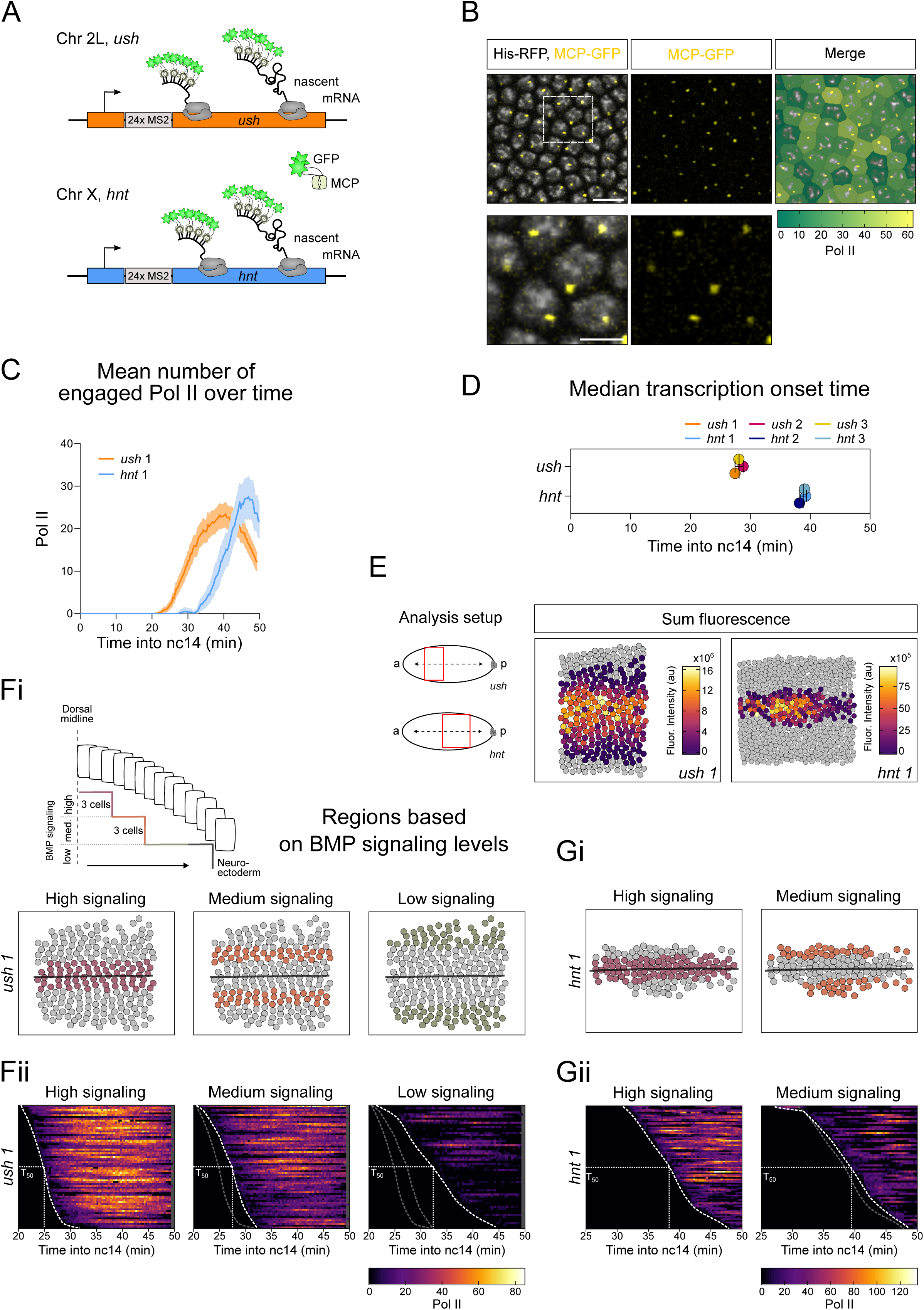
Temporal dynamics of BMP target gene transcription. (A) Schematic of the endogenous genomic locus for *ush* (orange) and *hnt* (blue) with 24xMS2 loops inserted into the 5’UTR. (B) Region from a still (Video S1) maximum projected showing active *ush* transcription by MCP-GFP fluorescence and converted to Pol II number. Enlarged region (lower) shows one active transcription site per nucleus. (C) Mean *ush* and *hnt* transcription traces over time for one representative embryo (209, *ush* and 144 *hnt* active nuclei). See Figure S1C, D for biological replicates. (D) Median transcription onset times of *ush* and *hnt* biological replicates (n = 3 for *ush* 209, 186, 223 nuclei and *hnt* 144, 171, 229 nuclei). (E) Schematics of the *ush* (anterior) and *hnt* (central) analysis domains. Cumulative expression domains of representative embryos, colored by sum fluorescence produced by nuclei throughout nc14 (note different scales). (F) Schematic summarizing the BMP gradient with cells receiving high, medium and low signaling. The *ush* expression domain of a representative embryo is false colored, highlighting the signaling regions, spanning 3, 3 and 4 cell rows respectively when mirrored at the dorsal midline (i). Heatmaps of single-cell traces from one representative embryo, sorted according to transcription onset (scale as indicated, grey indicates periods where nuclei were not tracked), n= 68 (high), 70 (medium), 60 (low) nuclei (ii). Time of transcriptional onset was traced to visualize onset fronts of different regions and the position at which half the nuclei in a region initiated transcription is indicated as T_50_ (25.16 min, 27.45 min and 33.5 min). (G) As in (F) showing the *hnt* expression domain of a representative embryo. As the *hnt* expression domain is very narrow, the high signaling region was defined to span 2 cell rows (i). Single cell traces for *hnt* transcription profiles (n = 83 and 61 nuclei, T_50_ = 43 min and 48.3 min) (ii). Note different scale to *ush*. Scale bar, 10 μm (B, top) and 3 μm (B, bottom). Mean ± 95% confidence intervals (C) or mean ± SD (D). **See also Figure S1, S2**.

Embryos were imaged prior to the onset of nuclear cleavage cycle 14 (nc14) to allow accurate timing of the initial activation of *ush* and *hnt* relative to the start of nc14 (Video S1 and S2 for *ush* and *hnt* transcription, respectively). We imaged the bulk of the expression domain for *ush*, whereas for *hnt* we imaged the central and posterior part; active nuclei are false colored in a still from the video (Fig. S1Bi). As the *ush* expression domain is largely uniform along the anterior-posterior (AP) axis, we have focused on the anterior part for subsequent analysis (Fig. S1Bii). *hnt* expression is more modulated along the AP axis (Ashe et al. 2000), therefore we have analyzed nuclei in the central region (Fig. S1Bii), corresponding to the presumptive amnioserosa. The mean transcription profiles of *ush* and *hnt* during nc14 are shown for representative embryos in Fig. 1C. We used smFISH data (see later, Fig. 4) to estimate the number of transcribing RNA polymerase II (Pol II) molecules based on the fluorescent signal.

**Figure 2:**
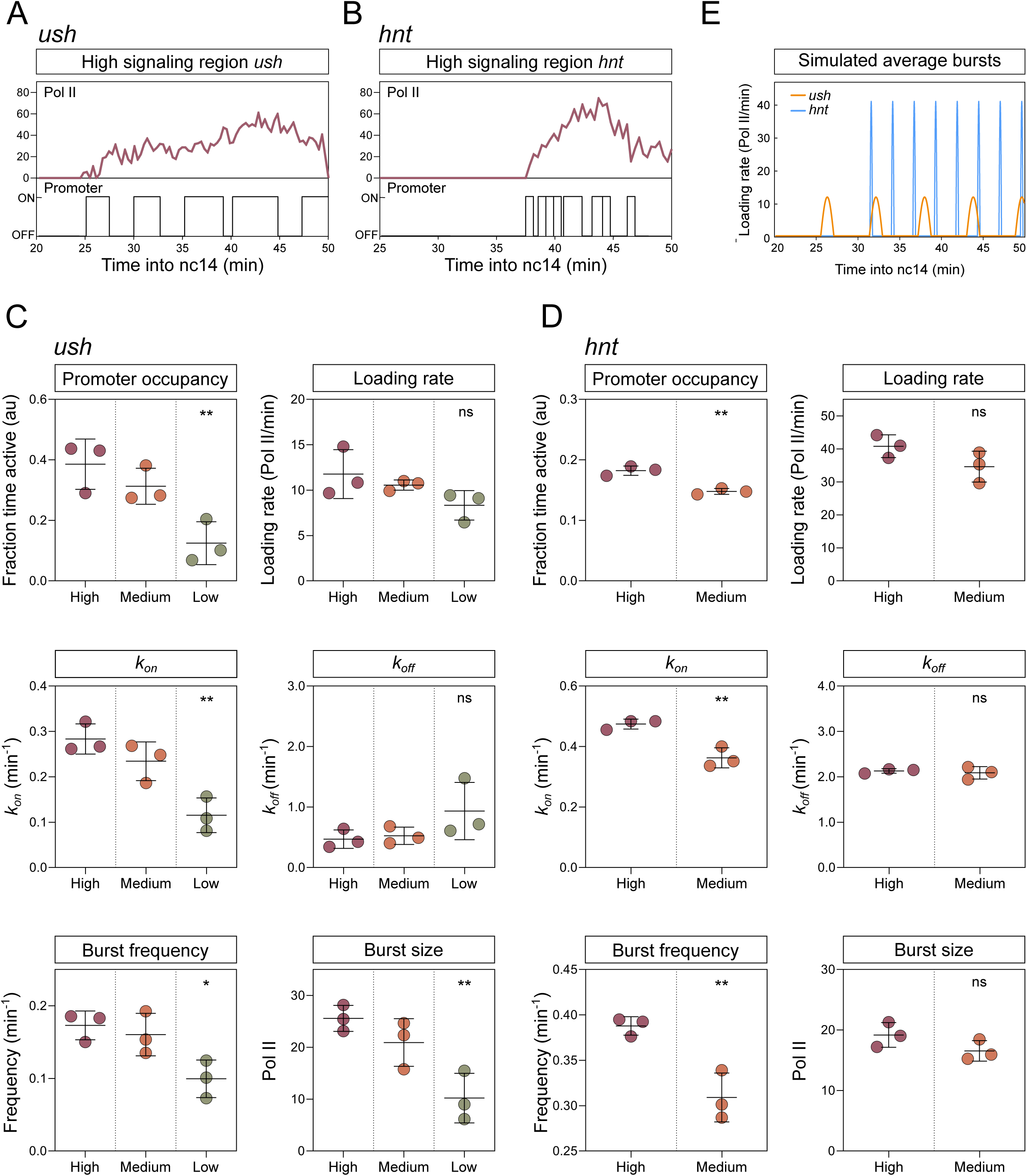
BMP signaling level modulates transcriptional burst kinetics. (A, B) Representative transcription traces from the high signaling region showing *ush* (A) and *hnt* (B) transcription and inferred promoter states. (C, D) Global analysis of burst parameters and rates, as labeled, for *ush* (C) and *hnt* (D) transcription in different spatial domains. Data points are colored according to BMP signaling regions. (E) Bursting simulation of *ush* (orange) and *hnt* (blue) transcription based on mean burst parameter values from the high signaling region and mean onset time of transcription. Mean ± SD (C, D) for n = 3 biological replicates. ^∗^p < 0.05, ^∗∗^p < 0.01, ns = not significant. One-way ANOVA with a Dunnett’s multiple comparisons test shows the difference to the high signaling region (C) or Student’s t-test (D). **See also Figure S3**.

**Figure 3:**
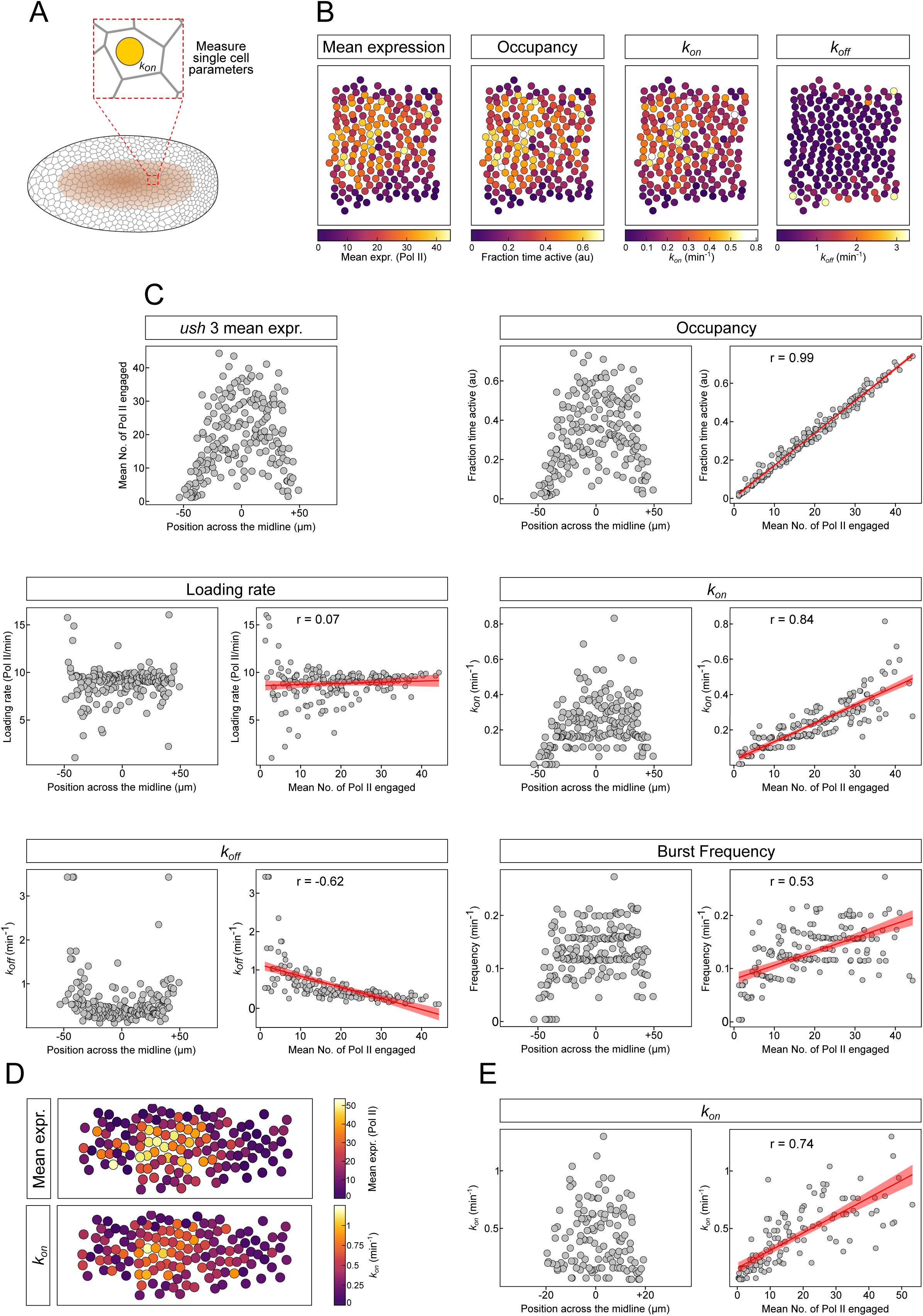
BMP signaling level is decoded by modulating *k*_*on*_. (A) Schematic of a *Drosophila* embryo indicating the *ush* expression domain and single cell analysis of transcription parameters. (B) Expression domain of a representative *ush* embryo shown as heatmaps with nuclei false colored according to their mean expression and single cell transcription parameters for occupancy, *k*_*on*_ and *k*_*off*_. (C) Mean expression values of *ush* plotted according to the nuclear position across the dorsal midline for one representative embryo. Burst kinetic parameters of single *ush* nuclei plotted based on dorsal midline position (left) and against mean expression (right) for promoter occupancy, loading rate, *k*_*on*_, *k*_*off*_ and burst frequency. See Figure S4A. (D) Heatmaps as in (B) showing mean expression and *k*_*on*_ for a representative *hnt* expression domain. (E) Single cell *k*_*on*_ values plotted based on their position (left) and against their mean expression (right). See Figure S4B-D for other *hnt* parameters. Pearson correlation coefficient is shown for each pair of variables between burst parameter and mean expression. Linear regression is shown ± 95% confidence intervals. n= 195 nuclei for *ush* (189 for loading rate) and 131 nuclei for *hnt* single cell parameters. **See also Figure S4**.

**Figure 4:**
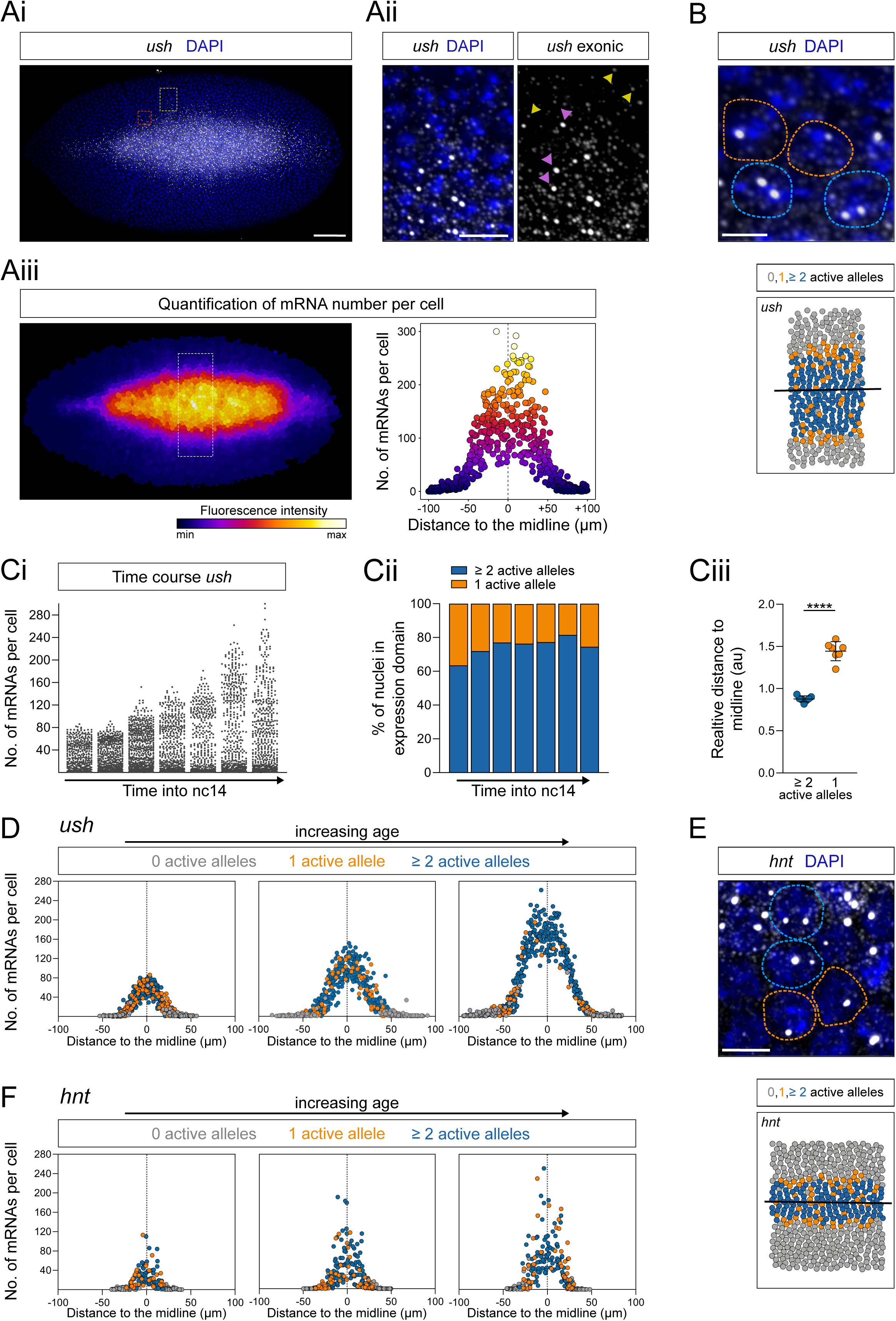
BMP target genes show stochastic transcription and graded mRNA outputs. (A) Fixed *Drosophila* embryo stained with exonic *ush* smFISH probes (grey) and DAPI (blue) (i). Inset outlined by the red box shown in (B) and by the yellow box in (ii) highlighting nascent transcription sites (pink arrowheads) and single mRNAs (yellow arrowheads). Image false colored by the combined fluorescence of individual mRNAs per cell (iii). Graph shows the quantification of the number of mRNAs per cell. Each point represents a nucleus from the domain outlined by the white rectangle in Aiii. (B) Enlarged region from (Ai) showing nuclei with one (orange outline) or two alleles (blue outline) actively transcribing *ush*. Schematic shows the expression region outlined in (Aiii) with nuclei false colored according to the number of active transcription foci. A line of best fit shows the middle of the expression domain. (C) Number of *ush* mRNAs per cell in embryos of increasing age in nc14 (i), the percentage of nuclei with one (orange) or two (blue) active alleles in each time course embryo (ii) and their median distance to the middle of the expression domain (iii) (n = 7 embryos). (D) Number of *ush* mRNAs per cell plotted according to the distance from the dorsal midline, at three time points (data from embryos 1, 4 and 6 in Ci). Embryos are ordered by increasing age and points are colored based on the number of active transcription foci. (E) As in (B) showing an enlarged region of an smFISH image indicating nuclei transcribing one (orange outline) or two *hnt* alleles (pink outline). Full image in Figure S5A. (F) as in (D) showing *hnt* time course data. Scale bar, 50 μm (Ai), 10 μm (Aii) or 5 μm (B, E). ^∗∗∗^p < 0.001. Mean ± SD (Ciii). Student’s t-test (Ciii). **See also Figure S5**.

Relative to *ush, hnt* has a similar number of transcribing Pol II (Fig. 1C, S1C, D) whereas both the transcription onset time, based on the first time a fluorescent signal is detected, and the time taken to reach maximal transcriptional activity are delayed (Fig. 1D, S1C, D). The transcription onset times for *ush* and *hnt* in each nucleus relative to its AP or dorsal-ventral (DV) position show little modulation along the AP axis (Fig. S1E). However, the onset times of *ush* and to a lesser extent that of *hnt* expression are delayed in nuclei further from the dorsal midline (Fig. S1F). The total transcriptional activity is found to be highest in nuclei closer to the dorsal midline, experiencing peak BMP signaling levels, for both *ush* and *hnt* (Fig. 1E). This difference in transcriptional activity depending on nuclear position is evident across developmental time (Fig. S1G).

While here we focus on embryos that are heterozygous for the MS2 engineered *ush* or *hnt* allele, analysis of MS2 homozygous embryos reveals that both alleles within each nucleus are similar in terms of fluorescent traces and onset times (Fig. S2A, B, Video S3). The difference between onset times increases in nuclei further from the dorsal midline, but is low overall, indicating similar times of activation for the alleles within a nucleus (Fig. S2B). Additionally, alleles in the same nucleus have more similar MS2 fluorescence than that of an allele in a neighboring nucleus or one randomly selected from within the expression domain (Fig. S2C). These data showing similar behavior of both alleles in a nucleus suggest that analysis of one is sufficient to capture representative transcriptional activity across the expression domain.

As the *hnt* and *ush* transcriptional activity varies between expressing nuclei, we performed K-means clustering analysis based on mean expression for all nuclei within the expression domain. For *ush* these data show that the nuclei partition into 3 clusters, which broadly map to the center, intermediate area and edges of the expression domain, whereas the narrower *hnt* expression domain is represented by 2 clusters (Fig. S1H). As the nuclei broadly partition into regions receiving different BMP signaling levels, for subsequent analysis we grouped nuclei based on position from the dorsal midline to more accurately relate the different groups to distinct signaling levels (Fig. 1Fi, Gi). Visualization of the individual fluorescent traces for all nuclei within these regions as heatmaps shows that nuclei from the middle of the expression domain receiving high signaling have a faster onset time and higher Pol II number than nuclei in the intermediate region with a further reduction in cells at the edge of the expression domain receiving low signaling levels (Fig. 1Fii). Similar findings are obtained for *hnt*, as nuclei receiving high Dpp signaling have faster onset times and more transcribing Pol II (Fig. Gii). These live imaging data show that cells vary in their transcriptional activity depending on their position with respect to the BMP gradient.

### Different BMP signaling levels alter transcriptional burst kinetics

Given the different transcriptional behaviors of nuclei, we further developed a previously described memory adjusted Hidden Markov Model (HMM) (Lammers et al., 2020) to infer rates and bursting parameters from the fluorescent traces, based on a two state promoter model (Fig. S3Ai. These include *k*_*on*_ and *k*_*off*_, the rates the promoter turns on and off; burst frequency; promoter occupancy, the fraction of time the promoter is active; *k*_*ini*_, Pol II loading rate; and burst size, which is the product of loading rate and duration (Zoller et al., 2018) (Fig. S3Aii). Using this computational approach, we can extract the promoter states corresponding to each time point in our live imaging data. Representative *ush* and *hnt* fluorescent traces for nuclei from the center of the expression domain receiving peak Dpp signaling and the inferred promoter states are shown in Fig. 2A, B, revealing different promoter activity profiles for the two Dpp target genes. Traces for nuclei at other positions in the expression domain are shown in Fig. S3B and C.

We grouped *ush* fluorescent traces based on nuclear position within the expression domain to infer the global kinetic parameters, with each group receiving progressively lower BMP signaling levels (as in Fig. 1Fi). For the narrower *hnt* expression domain, we separated cells into 2 regions (Fig. 1Gi). Expression is the product of promoter occupancy and Pol II loading rate, and as shown in Fig. 2C, D, for both *ush* and *hnt* there is a significant decrease in occupancy with lower BMP signaling levels, whereas the small reduction in loading rate is not significant. Therefore, the amount of signal does not alter the Pol II loading rate but instead modulates the fraction of time the promoter is active. The reduction in occupancy is through control of burst frequency and in particular *k*_*on*_, with both showing a significant reduction when signaling is lower (Fig. 2C, D). In contrast, *k*_*off*_ is unchanged in cells receiving different levels of signal (Fig. 2C, D), indicating that the duration of transcriptional bursts (equivalent to 1/*k*_*off*_) is not regulated. There is also a significant reduction in burst size for *ush*, but not *hnt*, when BMP levels decline (Fig. 2C, D). Based on the *ush* and *hnt* parameters the theoretical burst profiles of nuclei receiving peak Dpp signaling can be compared (Fig. 2E). These show that while *ush* is transcribed in relatively long duration bursts, *hnt* exhibits higher amplitude bursts of higher frequency and shorter duration (Fig. 2E). Despite these differences in amplitude and duration, burst size is similar in both cases, resulting in the synthesis of comparable numbers of *ush* and *hnt* mRNA in cells receiving peak signaling.

### Different Dpp concentrations control the rate the promoter switches on

This global analysis of cells separated into regions receiving different BMP levels shows that many parameters change, including *k*_*on*_, frequency, occupancy and in the case of *ush*, burst size. To investigate the cellular response in more detail and identify which parameter is the major determinant of the transcriptional response, we extended the modelling to infer burst parameters at single cell resolution (Fig. 3A, B). We then used these single cell parameters to assess the degree of correlation with the expression level for each nucleus (Fig. 3C). The fitted HMM model predicts “on” and “off” periods following a geometric distribution with parameters determined by *k*_*on*_ and *k*_*off*_ in each region and the single-cell analysis is consistent with this (Fig S3D). Promoter occupancy shows a very high correlation with mean expression, such that it almost perfectly predicts the expression level of every active nucleus (Fig. 3B, C). In contrast, Pol II loading rate is poorly correlated with expression (Fig. 3C, S4A), supporting the conclusion above that occupancy determines the response of cells to different levels of BMP signaling more than Pol II loading rate.

As promoter occupancy depends on *k*_*on*_ and *k*_*off*_, we next determined the contribution of each of these rates. As shown in Fig. 3C, *k*_*on*_ is strongly correlated with expression, unlike *k*_*off*_ (Fig. 3C, S4A), suggesting that promoter occupancy predicts expression predominantly through changes in *k*_*on*_. Consistent with *k*_*on*_ being important, burst frequency is also correlated with mean expression (Fig. 3C, S4A). Similar findings are obtained for *hnt* bursting parameters at single cell resolution (Fig. 3D, S4B), with promoter occupancy and *k*_*on*_ most correlated with mean fluorescence, followed by frequency, whereas loading rate and *k*_*off*_ is poorly correlated (Fig. 3E, S4C, D). Together these data provide further evidence that BMP signaling regulates target gene expression through modulating *k*_*on*_.

### Graded mRNA outputs in response to the BMP gradient

As *k*_*on*_ and transcriptional bursting for *ush* and *hnt* varies across the expression domain, we next investigated mRNA output. To this end, we quantitated the number of *ush* mRNAs using smFISH with probes targeting exonic sequences (Fig. 4Ai, ii). This analysis reveals a graded *ush* mRNA distribution, with lower numbers in cells further from the midline (Fig. 4Aiii), as predicted by a reduced rate of promoter activation in cells receiving lower BMP signaling. Also consistent with bursting, we find that our static snapshot smFISH images frequently only capture one transcriptionally active allele (Fig. 4B). Quantitation of the number of *ush* mRNAs from when the gene is first activated in nc14 embryos through developmental time reveals that the maximum number of transcripts per cell increases with age (Fig. 4Ci). Around one third to one quarter of the cells, mostly at the edge of the expression domain, had only one allele transcribing at the time the embryos were fixed, with a higher proportion when the gene first switches on (Fig. 4Cii-Ciii). The graded mRNA distribution in cells over time (Fig. 4D) mirrors that of the Dpp gradient, which starts shallow then refines (Dorfman and Shilo, 2001; Mizutani et al., 2005). While cells with one active allele are initially captured throughout the expression domain as the gene initiates, later in development they are more commonly found on the edge, where the low Dpp levels are limiting for promoter activation. As a result, cells near the midline accumulate ∼10-fold more mRNAs per cell compared to those at the edges of the expression domain (Fig. 4D).

Visualization and quantitation of *hnt* mRNAs using smFISH also revealed a gradient of mRNA output, with cells with a single active allele captured more frequently nearer the edge of the expression domain where Dpp signaling levels are lower (Fig. 4E, F, S5A, B). We also provide evidence for regulation of transcriptional bursting for *short gastrulation* (*sog*) and *brinker* (*brk*), target genes of the Dorsal gradient, depending on their position with respect to the gradient. For these target genes nuclei with a single active allele are predominantly detected on the edges of the expression domain (Fig. S5C-E). This is particularly evident on the dorsal border where activator is limiting, with such a bias not observed for Dpp target genes (Fig. S5F, G). Together, these data show graded mRNA distribution in response to the Dpp signaling gradient and, using detection of a single active allele as a proxy for bursting, suggest that bursting is spatially regulated during Dorsal gradient interpretation.

### The transcriptional response of *ush* to peak Dpp signaling is saturated

The above analyses suggest that higher Dpp signaling increases the rate with which the *ush* and *hnt* promoters become active, resulting in increased mRNA output. To address the effect of additional Dpp on bursting and mRNA numbers, firstly we imaged *ush* transcription in the presence of ectopic signaling by introducing a single copy of the st2-*dpp* transgene (Ashe et al., 2000) (Video S4). For the analysis we focused on cells in the region where st2*-dpp* is expressed, which have an expanded *ush* expression pattern (Fig. 5A), and the equivalent stripe 2 region in wt *ush* embryos. The *ush* transcription onset time is slightly earlier in the presence of st2-*dpp* (Fig. 5B) and the number of transcribing Pol II is increased, although the time at which maximum transcription is reached is similar to wt (Fig. 5C).

**Figure 5:**
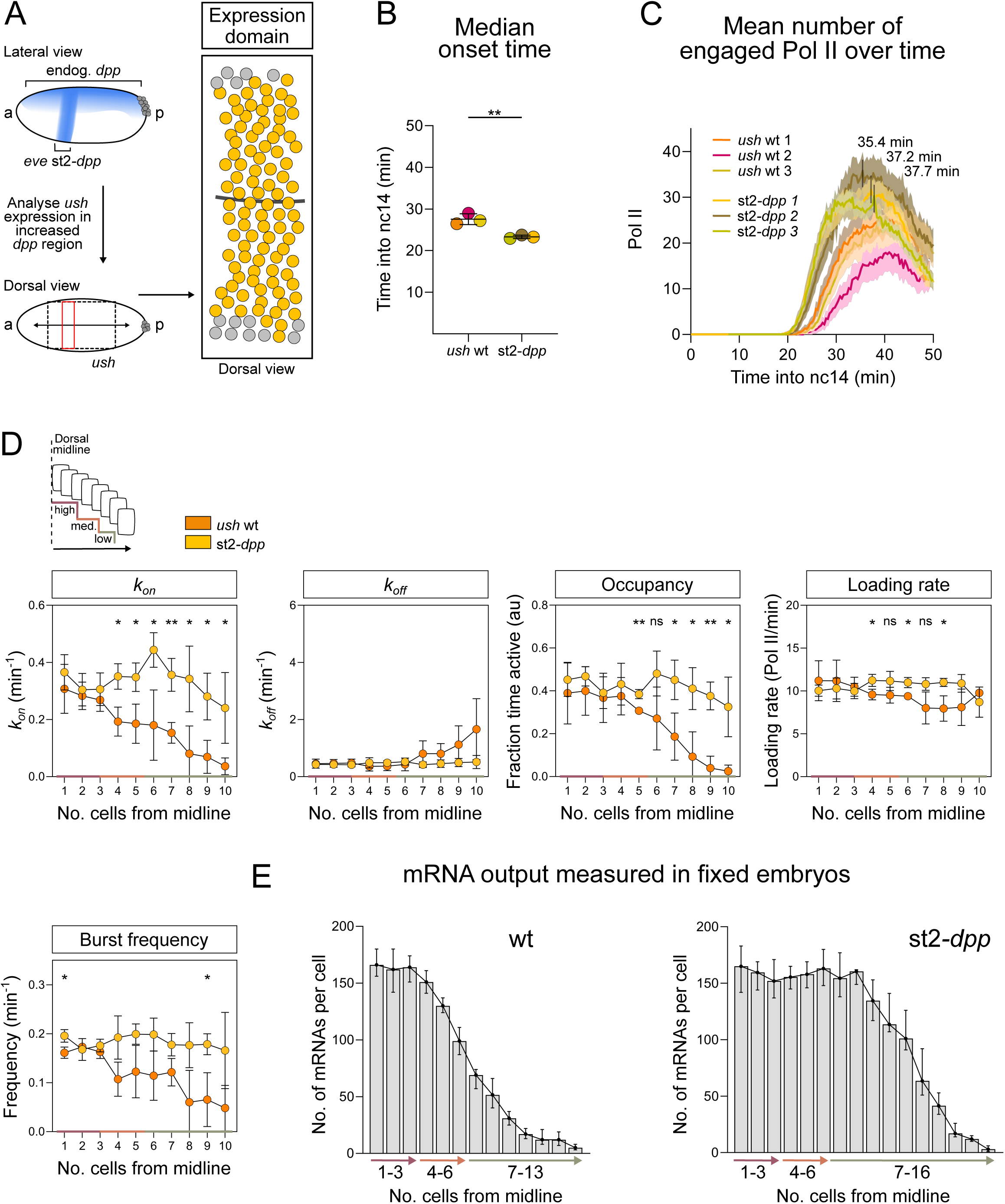
Increased BMP signaling increases the promoter activation rate. (A) Schematic showing ectopic *dpp* expression from the st2-*dpp* transgene, the imaging region (dotted box) and analysis domain (red box). The cumulative expression domain shows all active nuclei in one representative embryo. (B) Transcription onset times of *ush* shown for biological replicates in wt and st2*-dpp* embryos. (C) Mean *ush* transcription over time in st2*-dpp* and wt embryos. Maximum signal time point for st2*-dpp* is indicated. (D) Schematic indicates regions relating to BMP signaling levels. Single cell transcription parameters are plotted in 1 cell wide regions, based on the median parameter values for each biological replicate, n=3. The expanded st2-*dpp* expression (region 4) resulted in mean values for cell rows 11 and 12 as follows: *k*_*on*_ (0.13 min^-1^), *k*_*off*_ (0.66 min^-1^), promoter occupancy (0.22), loading rate (21.4 Pol II/min) and burst frequency (0.09 min^-1^). (E) Number of *ush* mRNAs per cell in fixed *Drosophila* embryos, reported in 1 cell wide bins from the dorsal midline (n= 780 wt, 297 st2-*dpp* nuclei). Mean ± SD (B, D), mean ± 95% confidence intervals (C) or median ± 95% confidence intervals (E), n= 94, 79, 113 nuclei for wt subsets to only include the st2 region and 121, 89, 107 nuclei for st2*-dpp* (B-D). ^∗^p < 0.05, ^∗∗^p < 0.01, ns = not significant. Student’s t-test (B, D).

We next used the memory adjusted HMM to infer burst parameters, after dividing the expression domain into 4 regions based on distance from the midline. The first three are equivalent to the regions described above for wt embryos (Fig. 1F), whereas the extra region reflects the expanded *ush* expression domain in the presence of st2-*dpp*. Note that we cannot analyze the full expansion of the *ush* expression domain as it extends too laterally to be captured accurately on our dorsally imaged embryos.

We inferred global and single cell parameters, then used the latter to plot the data in single cell wide bins moving out from the dorsal midline (Fig. 5D, see legend for data for region 4 for st2-*dpp* cells). These data reveal that in the first 3 cell rows mirrored at the dorsal midline, where there is maximal Dpp levels in a wildtype embryo (Dorfman and Shilo, 2001; Mizutani et al., 2005), addition of extra Dpp does not significantly alter the bursting kinetics. However, in cell widths 4-10 inclusive, which would normally receive medium and low Dpp, the ectopic Dpp significantly increases *k*_*on*_, with increases in occupancy and burst frequency observed also (Fig. 5D). In cell widths 4-10 inclusive, *k*_*on*_ reaches the rate observed in cells receiving peak Dpp signaling in wt embryos. As this rate does not increase in cells near the midline when extra Dpp is introduced, this suggests that *k*_*on*_ is already close to saturation in wildtype cells receiving the highest Dpp signaling. In contrast, *k*_*off*_ is unchanged and loading rate is only mildly affected in some cells when additional Dpp is introduced (Fig. 5D), consistent with the findings above that these parameters are not highly sensitive to alterations in Dpp concentration.

We also used smFISH to quantitate *ush* mRNA numbers in st2-*dpp* embryos, again focusing on the area where st2-*dpp* is expressed. This analysis shows that the mRNA number is unchanged in cells near the midline, but is increased in cells further away (Fig. 5E). As such, the drop in mRNA levels detected in wt embryos after row 4 is not observed (Fig. 5E), consistent with the Dpp increasing *k*_*on*_ in those cells (Fig. 5D). Together, these data further support the conclusion that Dpp signaling levels are decoded through changes in *k*_*on*_, resulting in a greater transcriptional output in cells normally receiving medium and low Dpp. These data also suggest that the *ush* transcriptional output is at its maximum in cells at the midline.

### Modulation of *k*_*on*_ by the enhancer and promoter

As the above data suggest that BMP signaling level predominantly regulates *k*_*on*_, we next addressed the role of the promoter in the transcriptional response by replacing the *ush* promoter with that of *hnt* in the endogenous locus (*hnt>ush*) (Fig. 6A). This line also contains 24 copies of the MS2 stem loops in the *ush* 5’UTR as described above so that the effect of changing the promoter on burst kinetics can be determined. Analysis of the fluorescent signals for *hnt>ush* (Video S5) reveals that the cumulative expression pattern, comprised of every cell that activates transcription at one or more time points, is similar but slightly narrower compared to wt *ush* (Fig. S6A). The times of transcription onset and at which maximum fluorescence is reached for *hnt>ush* are equivalent to those observed for *ush* (Fig. 6B, C, Fig. S6B). As *hnt* has a later onset time than *ush* (Fig. 1D) and changing the *ush* promoter to that of *hnt* has no effect on onset time (Fig. 6B), this suggests that onset time is largely dictated by the enhancer, with only fine-tuning by the promoter. The *hnt>ush* embryos analyzed have a small increase in the number of transcribing Pol II molecules (Fig. 6C, S6B).

**Figure 6:**
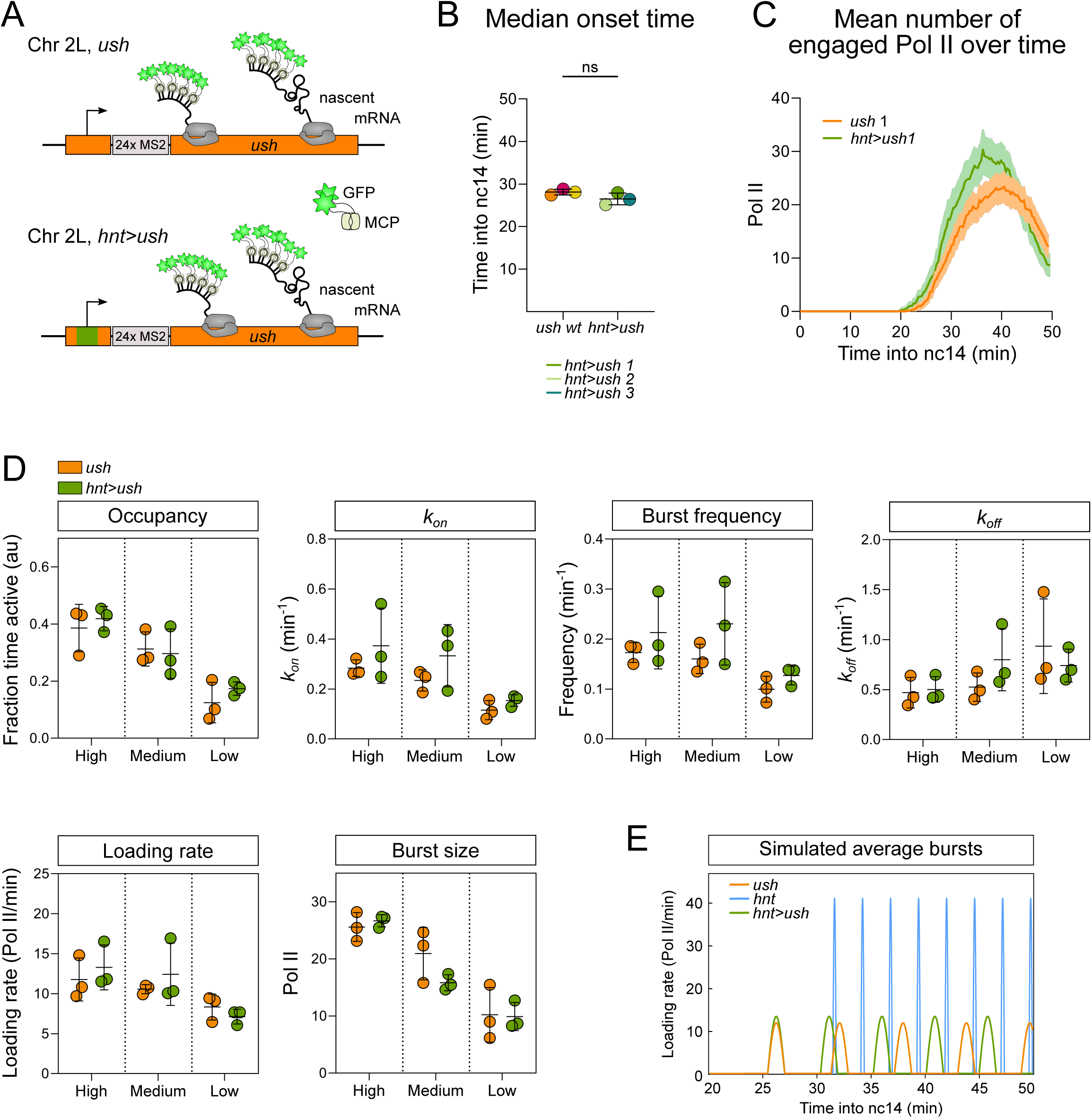
Insertion of a heterologous promoter does not alter Pol II loading rate. (A) Summary of the endogenous *ush* locus with its own promoter (orange) or the *hnt* promoter (green) and 24xMS2 loops present in the 5’ UTR. (B) Median onset times of *ush* transcription in *ush* and *hnt>ush* from n= 3 biological replicates with 209, 186, 233 and 154, 187, 202 nuclei for *ush* and *hnt>ush* respectively. (C) Mean transcription of *ush* in representative embryos with 209 (*ush*) and 154 (*hnt>ush*) active nuclei. See Figure S6B. (C) Global analysis comparing burst parameters for *ush* transcription spatially and between genotypes. (E) Bursting simulation of *ush* (orange), *hnt* (blue) and *hnt>ush* (green) transcription based on mean burst parameter values from the high signaling expression domains and typical transcription onset times. Mean ± SD (B, D) or mean ± 95% confidence intervals (C). Student’s t-test (B) between genotypes in each region (D). **See also Figure S6**.

Cells expressing *hnt>ush* were subdivided into 3 regions (Fig. S6Ci), as described for wt *ush*. Within the high and medium regions, the fluorescent traces show a small increase in transcribing Pol II (Fig. S6Cii) that is not statistically significant when compared between all biological replicates (Fig. S6Ciii). We used these 3 regions to infer global bursting parameters from the model (Fig. 6D). The global parameters show that there is no significant change in any of the parameters when the *ush* promoter is replaced with that of *hnt* (Fig. 6D). Comparison of the *hnt>ush* burst profile with that of wt *ush* and *hnt* in cells receiving peak signaling, based on the inferred parameters, shows a greater similarity to that of wt *ush* (Fig. 6E). There are modest increases in mean *k*_*on*_, frequency and loading rate for *hnt>ush* compared to wt *ush*, which likely underpins the small increase in the number of transcribing Pol II detected (Fig. 6C, S6C). Additionally, *k*_*on*_ and frequency are more variable between *hnt>ush* replicate embryos (Fig. 6D). Pol II loading rate is higher for *hnt* than *ush* (Fig. 2C) and the *hnt>ush* loading rate is typical of *ush* (Fig. 6D, E), suggesting that it is dictated by the enhancer. The data described above show that *k*_*on*_ is the rate that is sensitive to BMP levels and, as this does not change significantly in the presence of the *hnt* promoter, it also appears that in these embryos *k*_*on*_ is predominantly dictated by the enhancer.

We also analyzed *ush*>*hnt* embryos with the reciprocal promoter swap, which have a similar expression pattern to that of wt *hnt* (Fig. 7A, S7A, Video S6). However, the mean number of Pol II transcribing *ush*>*hnt* is reduced relative to *hnt* (Fig. 7B, S7B, C). The onset time, the time transcription is first detected in any nucleus within the expression domain, is not significantly different for *ush*>*hnt* (Fig. 7C), although two embryos show a delay. A short delay in *ush*>*hnt* mean onset time is apparent when nuclei are separated into the two signaling regions (Fig. S7C). Analysis of the global parameters for the two *ush*>*hnt* regions (Fig. 7D) reveals that the *ush*>*hnt* burst profile is more similar to *hnt* than *ush* (Fig. 7E). However, *ush*>*hnt* has a significantly reduced promoter occupancy, compared to *hnt* embryos, which appears to be driven primarily by a reduction in *k*_*on*_ in both regions, although *k*_*off*_ is also increased in cells receiving medium signaling (Fig. 7D). Consistent with this, burst frequency is reduced. The similar mean loading rate for *ush*>*hnt* to that of *hnt*, again consistent with it being dictated by the enhancer. However, loading rate is highly variable between *ush*>*hnt* embryos, as is therefore burst size (Fig. 7D), potentially suggesting some incompatibility between the enhancer and promoter (see Discussion). The reduction in *k*_*on*_ and burst frequency in *ush*>*hnt* embryos when the only sequence changed is the core promoter, suggest that the promoter sequence itself impacts on the ability of the enhancer to promote its activation and therefore the frequency of transcriptional bursts.

**Figure 7:**
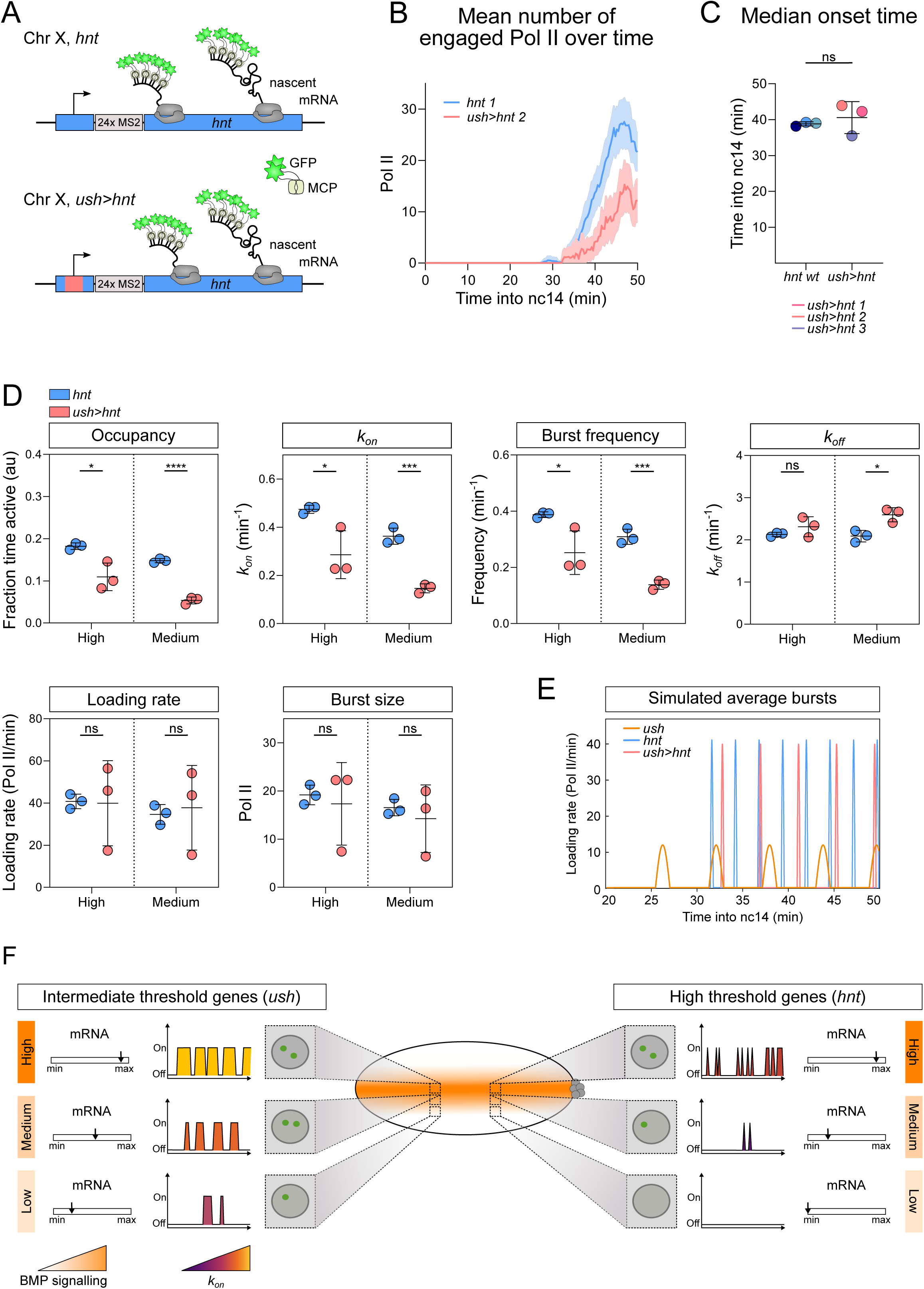
Replacement of the *hnt* promoter reduces burst frequency. (A) Schematic of the *hnt* locus with its own promoter (blue) or the *ush* promoter (pink) and 24xMS2 loops in the 5’ UTR. (B) Mean *hnt* transcription output in representative *hnt* (144 nuclei) and *ush>hnt* (133 nuclei) embryos. See Figure S7B. (C) Median onset times of *hnt* transcription in *hnt* and *ush>hnt* embryos with n=3 biological replicates with 144, 171, 229 and 133, 187, 202 nuclei for *hnt* and *ush>hnt* respectively. (D) Comparison of global burst parameters for *hnt* transcription spatially and between genotypes. Data points are colored according to genotype. (E) Simulation of average transcriptional bursts of *hnt* (blue), *ush* (orange) and *ush>hnt* (pink) based on mean burst parameter values from the high BMP signaling region and typical transcription onset times. (F) Summary of the promoter state profiles for *ush* and *hnt*, how these respond to altered BMP signaling levels, and the resulting effects on allele activity and mRNA number. Mean ± 95% confidence intervals (B) or mean ± SD (C, D). ^∗^p < 0.05, ^∗∗∗^p < 0.001, ^∗∗∗∗^p < 0.0001, ns = not significant Student’s t-test (C) between genotypes in each region (D). **See also Figure S7**.

## DISCUSSION

Here we analyze the transcriptional burst kinetics of the endogenous *hnt* and *ush* genes at single cell resolution. We show that cells interpret different BMP signaling levels by modulating burst frequency via *k*_*on*_ (Fig. 7F). For *ush, k*_*on*_ is unchanged when the *hnt* promoter is introduced, suggesting that features of the enhancer, most likely transcription factor binding sites, dictate *k*_*on*_. Transcription factor binding is coupled to initiation of a burst (Donovan et al., 2019), and burst frequency depends on the time it takes a transcription factor to find its binding site (Larson et al., 2011), providing an explanation for why high BMP/pMad levels increase *k*_*on*_. Consistent with this, other studies have found transcription factor concentration and enhancer strength to regulate *k*_*on*_ (Fukaya et al., 2016; Lammers et al., 2020; Larson et al., 2013; Larsson et al., 2019; Senecal et al., 2014). For *ush*, we provide evidence that *k*_*on*_ is at a maximum in cells receiving peak signaling. A burst frequency ceiling has been described for TNF targets (Dar et al., 2012), although the *k*_*on*_ ceiling for *ush* appears to be gene specific, as it is below that observed for *hnt* and other *Drosophila* genes studied (Lammers et al., 2020; Zoller et al., 2018). Perhaps *k*_*on*_ for *ush* is limited by slow recruitment of pMad or fewer productive activation events between its enhancer and promoter. Current ideas for enhancer-promoter communication include a dynamic kissing model that invokes transient interactions (Cho et al., 2018) or models based on the proximity of regulatory elements in space (Andersson and Sandelin, 2020; Heist et al., 2019), potentially in phase separated condensates (Hnisz et al., 2017). In these models, the Mediator coactivator may act as a dynamic molecular bridge between the enhancer and promoter, as activator-recruited Mediator at the enhancer can also make contact with the promoter (Soutourina, 2019).

When the *hnt* and *ush* promoters are swapped, mean Pol II loading rate is unchanged in both cases suggesting that it is predominantly dictated by the enhancer, although we detect increased variability in loading rate for *ush>hnt*. This suggests that transcription factors bound to the enhancer control Pol II loading rate, which is consistent with a previous study that found loading rate to be influenced by the strength of the transcription factor’s activation domain (Senecal et al., 2014). Loading rate is unchanged by altered BMP levels, with a constant loading rate also described for bursting of gap genes (Zoller et al., 2018), suggesting it is not a major regulated step of transcription in the *Drosophila* embryo. Despite the similar bursting behavior between *ush* and *hnt>ush*, introducing the *ush* promoter into *hnt* reduces *k*_*on*_. This suggests that the *hnt* promoter has some feature that is important in the *hnt* genomic context, for example a TATA box or highly paused Pol II, as the *ush* promoter lacks both (Saunders et al., 2013). TATA increases burst frequency in the presence of interferon signaling (Hendy et al., 2017), however other studies have linked TATA promoters to burst size (Hornung et al., 2012; Larsson et al., 2019) and the *hnt* promoter does not increase frequency when introduced into *ush*. Another possibility is that the ability of paused Pol II at the *hnt* promoter to prevent encroachment of nucleosomes (Gilchrist et al., 2010) increases *k*_*on*_. In support of this, introduction of nucleosome disfavoring sequences around a promoter and linked transcription factor binding site was found to increase burst frequency (Dadiani et al., 2013). It has also been shown that forcing interactions between a β-globin enhancer and its promoter increases burst frequency (Bartman et al., 2019). As enhancer-promoter compatibility has been proposed (Catarino and Stark, 2018), perhaps the *hnt* enhancer is more compatible with its cognate promoter. In terms of compatibility, Mediator and the p300 acetyltransferase, both of which are recruited by Smads (Hill, 2016), most strongly activate TATA containing promoters (Haberle et al., 2019). While further work is required to understand how the core promoter influences bursting, our results suggest a role for both the enhancer and promoter in influencing *k*_*on*_ and burst frequency, thereby allowing greater flexibility in controlling the transcriptional response.

We show that *hnt* transcriptional bursts are shorter and of higher frequency and amplitude than the *ush* bursts. The resulting *hnt* promoter occupancy is around half that for *ush*, providing a molecular explanation for the observed threshold responses of these genes to the BMP gradient (Fig. 7F). Unlike for *ush*, low BMP signaling levels are insufficient to maintain the *hnt* promoter in an active state, resulting in a narrow expression pattern. Burst duration, although not responsive to Dpp levels, is around 4 times longer for *ush* than for *hnt*. Transcription factor dwell time, which is limited by binding site affinity and nucleosomes, controls burst duration (Donovan et al., 2019; Senecal et al., 2014). As the Smad proteins have low affinity for DNA and weak specificity, they cooperate with other DNA binding proteins (Hill, 2016). The *ush* and *hnt* enhancers have yet to be characterized, but the pioneer factor Zelda and homeodomain protein Zerknüllt may be pMad cofactors for intermediate (*ush*) and peak (*hnt*) Dpp targets, respectively (Deignan et al., 2016; Xu et al., 2005) It is possible that at the *hnt* enhancer the Zerknüllt-pMad complex has a shorter residency time or the pMad binding sites are weaker affinity, resulting in shorter duration bursts.

The lack of a contribution of burst duration (1/*k*_*off*_) to decoding BMP signaling is in stark contrast to the findings that Notch alters the duration, but not frequency, of transcription bursts in *Drosophila* and *C. elegans* (Falo-Sanjuan et al., 2019; Lee et al., 2019). Increasing gene expression through high *k*_*on*_ rates can decrease noise, whereas lengthening burst duration is associated with more noise (Wong et al., 2018). Regulation of burst frequency may also allow genes to respond more sensitively to activator concentration (Li et al., 2018). Therefore, perhaps regulation of BMP target genes via *k*_*on*_ has the advantage of allowing more sensitive regulation with less noise. It remains to be determined whether other signals will be interpreted through changes in *k*_*on*_ and burst frequency or duration.

The different burst kinetics of BMP target gene transcription across the expression domain explain why cells at the edge are frequently captured with a single active allele in the smFISH data. *sog* and *brk* exhibit transcriptional bursting (Esposito et al., 2016), and our data suggest that *sog* and *brk* bursting is regulated across their expression domains. Allele by allele repression has been observed in the *Drosophila* embryo, potentially because repressors are better able to act in the refractory period following a burst (Esposito et al., 2016). Such allele by allele repression can also explain why nuclei with one active allele are observed at the ventral borders of the *sog* and *brk* expression domains, where Dorsal activator levels are high. Single allele transcription has also been reported for zygotic *hunchback* (*hb*) transcription, which is activated by the Bicoid gradient, particularly at the borders of the expression domain (Lucas et al., 2013; Porcher et al., 2010). We suggest that infrequent transcriptional bursting, with a concomitant reduction in mRNA number, is a general feature of gradient interpretation for cells receiving low signal.

The *ush* mRNA distribution reflects the spatial BMP gradient as the central 6 rows that receive peak BMP signaling have the highest mRNA number/cell, with subsequent declining mRNA numbers mirroring a reduction in Dpp levels (Dorfman and Shilo, 2001; Mizutani et al., 2005). Additionally, modelling suggests that the concentration of BMP-receptor complexes at the dorsal midline doubles between 20 min and 30 min into nc14 (Mizutani et al., 2005; Umulis et al., 2006). This corresponds to the onset times of *ush* and *hnt*, respectively, suggesting that *hnt* transcription requires more activated receptors. Furthermore, BMP-receptor levels peak at ∼45 min into nc14 (Umulis et al., 2006), which broadly coincides with the observed maximum fluorescence output we detect for *ush* and *hnt*.

We suggest that altered transcriptional burst kinetics and graded mRNA numbers in response to morphogen gradients can impact on cell fate decisions. We propose that cells on the edge of an expression domain synthesize sufficient mRNAs to adopt a particular cell fate, whereas cells in the center have a surplus of transcripts. This model can explain the lack of robustness when shadow enhancers are deleted (Antosova et al., 2016; Frankel et al., 2010; Perry et al., 2010). Perturbation of the system, such as removal of a shadow enhancer, would lead to a further reduction in mRNA number per cell so that those on the edge would only just exceed the threshold level. Another challenge, such as high temperature or reduced activator level, would further decrease the transcriptional output such that there are insufficient mRNAs to specify the correct cell fate. It will be interesting in the future to test how the different numbers of mRNAs per cell from key BMP target genes impact on the robustness of dorsal ectoderm cell fate decisions.

## ACKNOWLEDGEMENTS

We thank Lauren Forbes-Beadle for helpful discussions, Bloomington *Drosophila* Stock Center for flies, JP Vincent lab for plasmids, Cambridge Fly Facility for microinjections, Peter March and Egor Zindy for image analysis advice and the University of Manchester Bioimaging Facility for support. This project was supported by a Wellcome Trust Investigator Award to H.L.A. and M.R. (204832/Z/16/Z) and Wellcome Trust PhD studentships to C.H. (205975/Z/17/Z), J.B. (215187/Z/19/Z) and T.G.M. (110566/Z/15/Z).

## AUTHOR CONTRIBUTIONS

Conceptualization: C.H., J.B., M.R., H.L.A.; Investigation: C.H, C.S., P.U.; Software: J.B., T.G.M.; Writing-original draft: C.H., H.L.A.; Writing–Reviews & Editing: J.B, T.G.M, C.S., P.U., M.R.; Supervision: M.R., H.L.A.; Funding Acquisition: M.R., H.L.A.

## STAR METHODS

### LEAD CONTACT AND MATERIALS AVAILABILITY

Further information and requests for resources and reagents should be directed to and will be fulfilled by the Lead Contact, Hilary L. Ashe. The custom analysis codes generated during this study to link Imaris output files are available at https://github.com/TMinchington/sass. The modelling software used in this study will be published elsewhere but is available from the corresponding author on request. Plasmids and fly lines generated in this study are available from the corresponding author on request.

### EXPERIMENTAL MODEL AND SUBJECT DETAILS

#### Experimental Animals

*Drosophila melanogaster* flies were grown and maintained at 18°C while fly crosses for imaging were raised and maintained at 25°C. All flies were raised on standard fly food (yeast 50g/L, glucose 78g/L, maize flour 72g/L, agar 8g/L, nipagen 27ml/L, and propionic acid 3ml/L). Embryos were collected on apple juice agar plates that contained yeast paste.

The following fly lines were used for experiments in this study; st2-*dpp* (Ashe et al., 2000), *y*^*1*^*w*;P{His2Av-mRFP1}II*.*2; P{nos-MCP*.*EGFP}2* (BDSC Stock# 60340, RRID:BDSC_60340), *y*^*1*^*w*^*67c23*^; *24xMS2-ush* (this study), *y*^*1*^*w*^*67c23*^; *hnt>24xMS2-ush* (this study), *y*^*1*^*M{vas-Cas9}ZH-2Aw*^118^, *24xMS2-hnt* (this study), *y*^*1*^*M{vas-Cas9}ZH-2Aw*^118^, *ush>24xMS2-hnt* (this study) and *y*^*1*^*w*^*67c23*^, which we used as wildtype.

#### Generation of endogenous MS2 lines

Live imaging fly lines were generated through a two-step method of CRISPR/Cas9 genome editing with homologous recombination and ϕC31 integrase-mediated site-specific transgenesis. All MS2 tagged lines generated for this study are homozygous viable and fertile.

First deletions in the 5’UTR regions of *ush* isoform RC (456 bps; Chr 2L: 523446-523902, dm6 genome) and *hnt* isoforms RA and RB (705 bps; ChrX: 4617319-4618023, dm6 genome) were generated. Two PAM sites (flyCRISPR Optimal Target Finder tool: http://flycrispr.molbio.wisc.edu/tools) were used to create double strand breaks.

The plasmid pTV^cherry^ (gift from the Vincent lab; DGRC #1338) was used as a donor plasmid containing an attP reintegration site flanked on either side by homology arm sequences. Homology arms were inserted using KpnI and SpeI restriction sites, respectively.

*<u>ush</u>* <u>HA1:</u> forward primer GGTACCgtgcatagccacgacgttagg, reverse primer GGTACCccggggacgagacgagacctctta

*<u>ush</u>* <u>HA2:</u> forward primer ACTAGTggaagtgacaacataattgcc, reverse primer ACTAGTtccaagccttcactccactc

*<u>hnt</u>* <u>HA1:</u> forward primer GCTAGCgaagggttgctggtcacc, reverse primer GCTAGCcattgggtgcgtgtgtgtg

*<u>hnt</u>* <u>HA2:</u> forward primer ACTAGTcaactgttgaacacaatttcac, reverse primer ACTAGTcacacatgcatacatccagtc

The pU6-BbsI-chiRNA plasmid (RRID:Addgene_45946) was used to deliver guide RNAs (gRNA). 5’ phosphorylated oligonucleotides were annealed and ligated into the BbsI restriction site. Together, gRNA plasmids and the donor plasmid were injected into Cas9 expressing flies (BDSC Cat# 51323, RRID:BDSC_51323) by the Cambridge University injection service.

*<u>ush</u>* <u>gRNA1:</u> forward primer cttcgtctcgtctcgtccccgctc, reverse primer aaacgagcggggacgagacgagac

*<u>ush</u>* <u>gRNA2:</u> forward primer cttcgattatgttgtcacttcccgt, reverse primer aaacacgggaagtgacaacataatc

*<u>hnt</u>* <u>gRNA1:</u> forward primer cttcgcgcaaataggattacacat, reverse primer aaacatgtgtaatcctatttgcgc

*<u>hnt</u>* <u>gRNA2</u>: forward primer cttcgattgtgttcaacagttgcga, reverse primer aaactcgcaactgttgaacacaatc

Next, the attB-attP system was used for site-specific reintegration. Reintegration fragments were inserted into the RIV^cherry^ plasmid (gift from the Vincent lab; DGRC #1331). Wildtype sequences of promoter and 5’UTR regions, previously removed in the CRISPR process, were inserted into RIV^cherry^ using the NotI site to reconstitute wildtype loci. The 24xMS2-loop cassette (pCR4-24xMS2L-stable, RRID:Addgene_31865) was inserted using the BglII site. The RIV^cherry^ plasmid was co-injected with a ϕC31 integrase plasmid (Injection service) into the balanced CRISPR fly lines. Successful transformants were balanced and the marker region was removed by crossing to a cre-recombinase expressing fly line (BDSC Cat# 1501, RRID:BDSC_1501).

#### Promoter swap fly lines *hnt>ush* and *ush>hnt*

To generate the *hnt>ush* fly line, the core promoter sequence of *hnt* was inserted into the previously generated fly line carrying the *ush* 5’UTR deletion and an attP site. The core *hnt* promoter sequence and annotation (200 bps; Chr X: 4,617,464 - 4,617,663 dm6 genome) was determined based on peaks from Global Run-On Sequencing (GRO-Seq) data (Saunders et al., 2013). After co-injection with the ϕC31 plasmid, successful transformants were crossed to a Cre-recombinase expressing fly line (BDSC Cat# 1501, RRID:BDSC_1501). The *ush>hnt* fly line was generated in the same manner by inserting the *ush* core promoter sequence (200 bps; Chr 2L: 523,636 - 523,835 dm6 genome) into the previously generated fly line carrying the *hnt* 5’UTR deletion and attP site, as described above. Full cloning details available upon request.

#### Fly strains and genetics

To observe transcription from our reporter MS2 lines, female flies of the genotype His2av-RFP; MCP-eGFP, also with one copy of the *st2-dpp* transgene (Ashe et al., 2000) as appropriate, were mated with males homozygous for the 24*xMS2 tagged target gene locus*. Embryos from this cross were used for live imaging experiments.

To observe transcription from two MS2 tagged *ush* alleles, female flies of the genotype His2av-RFP/*24xMS2-ush*; MCP-eGFP were crossed to *24xMS2-ush* males. The resulting embryos were imaged live and the data were screened and retained if two active transcription sites were visible.

## METHOD DETAILS

### In situ hybridization

Embryo collections (2-4h), RNA probe synthesis and in situ hybridization with digoxygenin-UTP-labelled (Sigma, 11277073910) or biotin-UTP-labelled probes (Sigma, 11685597910) were performed as described by (Kosman et al., 2004). Antisense probes were approximately 1kb in length (Primer sequences available upon request). The following primary and secondary antibodies were used: Sheep Anti-Digoxigenin Fab fragments Antibody, AP Conjugated (1:250 Roche Cat# 11093274910, RRID:AB_514497), mouse anti-biotin (1:250 Roche, 1297597), donkey anti-Sheep IgG Secondary Antibody, Alexa Fluor or 555 (1:500; Thermo Fisher Scientific Cat# A-21436, RRID:AB_2535857) and donkey anti-Mouse IgG Secondary Antibody, Alexa Fluor 647 (1:500; Thermo Fisher Scientific Cat# A-31571, RRID:AB_162542). Samples were incubated with DAPI (1:500; NEB, 4083) and mounted in ProLong™Diamond Antifade Mountant (Thermo Fisher, P36961). For alkaline phosphatase blue stain; embryos were incubated in staining solution containing 0.675mg/mL nitro blue tetrazolium (NBT Sigma Cat# 11585029001) and 0.35mg/mL 5-bromo-4-chloro-3-indolyl-phosphate (BCIP Sigma Cat# B6149). Embryos were mounted in Permount™(BioWORLD Cat# 21750009).

### DNA oligonucleotides

Exonic probe sets for smFISH (Biosearch Technologies) were conjugated to Quasar 570 fluorophores (Table S1).

### smFISH

Fixed 2-4h old embryos were transferred into Wheaton vials (Z188700-1PAK, Sigma), washed 5 min in 50% methanol/50% phosphate-buffered saline with 0.1% Tween-20 (9005-64-5, Sigma) (PBT), followed by four 10 min washes in PBT, a 10 min wash in 50% PBT/5% wash buffer (10% formamide in 2X SSC; 300mM NaCl and 30mM trisodium citrate adjusted to pH 7) and two 5 min washes in 100% wash buffer. Next, embryos were incubated 2h at 37°C in smFISH hybridization buffer (2.5mM dextran sulphate, 10% formamide in 2X SSC). Probes were diluted in hybridization buffer to a final concentration of 1.25mM for smFISH Stellaris probes. Embryos were incubated in probe solution for 14h at 37°C, washed 15 min in pre-warmed hybridization buffer at 37°C, followed by three 15 min washes in pre-warmed wash buffer at 37°C. At room temperature, embryos were washed for 15 min in wash buffer and three times for 15 min in PBT in the dark. Embryos were blocked in 1xWestern Blocking Reagent (Sigma Cat# 11921673001) for 30 min, followed by incubation with anti-Spectrin (1:50 DSHB Cat# 3A9 (323 or M10-2)) overnight at 4°C in the dark. Embryos were then washed three times in PBT in the dark. One of the PBT washes included DAPI (1:500). Embryos were then mounted in ProLong Diamond Antifade Mountant (Thermo Fisher Scientific, Cat# P36961). All washes were performed with agitation.

### FISH/smFISH microscopy

Images were acquired with a Leica TCS SP8 AOBS inverted microscope using a 40x/ 1.3 HC Pl Apo CS2 or 63x/ 1.4 Plan APO objective with 8x or 2x line averaging, respectively. The confocal settings were as follows, pinhole 1 airy unit, scan speed 400Hz unidirectional line scanning and a format of 2048 x 2048 pixels. Images were collected with either Photon Multiplying Tube Detectors or Hybrid Detectors and illuminated using a white laser. The following detection mirror settings were used: Photon Multiplying Tube Detector DAPI excitation at 405nm (8%, collection: 415-474nm); Hybrid Detectors: AlexaFluor 488 excitation at 490nm (10%, 1 to 6us gating, collection: 498-548nm) and Quasar 570 excitation at 548nm (21%, 1 to 6us gating, collection: 558-640nm). All images were collected sequentially and optical stacks were acquired at 200nm spacing. Raw images were then deconvolved using Huygens Professional software (SVI, RRID:SCR_014237) and maximum intensity projections are shown in the figures. Unless stated otherwise in the figure legends, all embryos are oriented dorsally with the anterior to the left.

### Live Imaging microscopy

Embryos were dechorionated in bleach and positioned dorsally on top of a coverslip (Nr. 1, 18x 18 mm; Scientific Laboratory Supplies, Cat#MIC3110), thinly coated with heptane glue. A drop of halocarbon oil mix (4:1, halocarbon oil 700: halocarbon oil 27; Sigma Cat# H8898 and Cat# H8773) was placed in the middle of a Lumox imaging dish (Sarstedt AG & Co, Cat#94.6077.305) and two coverslips (Nr. 0, 18x 18mm; Scientific Laboratory Supplies, Cat#MIC3100) were placed on either side of the oil drop, creating a bridge. The coverslip with the embryos glued to it was then inverted into the oil, sandwiching the embryos between the imaging dish membrane and the coverslip.

Embryos were imaged on a Leica TCS SP8 AOBS inverted confocal microscope with a resonant scan head, using a 40x/1.3 HC PL apochromatic oil objective. Images of embryos heterozygous for MS2 imaging loci were obtained with the following confocal settings, pinhole 1.3 airy units, scan speed 8000Hz bidirectional, format 1024 x 700 pixels at 8 bit. Images were collected using hybrid detectors and the white laser at 70% with 488nm (8%) and 574nm (2%) at 8x line averaging for *ush* and *hnt>ush* embryos, with 488nm (25%) and 574nm (2%) at 8x line averaging for *hnt* and with 488nm (30%) and 574nm (2%) at 8x line averaging for *ush>hnt* embryos. Three-dimensional optical sections were acquired at 1 µm distance, a final depth of 55 µm and a final temporal resolution of 20 seconds per time frame. Images of embryos homozygous for the *ush* imaging locus were obtained using the microscope setup described above with the following alterations. Images were obtained at a 2.15 x confocal zoom and optical sections were obtained at 1μm with a final depth of 40μm and a final temporal resolution of 15.2 sec.

Images were processed with the Leica lightning deconvolution software. The mounting medium refractive index was estimated to be 1.41. Maximum intensity projections of 3D stacks are shown in the result sections. Embryos were imaged for 70-90 min and included the cleavage cycle of nc14 and the onset of gastrulation. During analysis all datasets were adjusted in time to account for slight temperature differences during imaging that can alter the speed of development. Therefore, nc14 was defined as the time between telophase of cleavage cycle 14 and the beginning of cephalic furrow formation. For the purpose of this study, nc14 was defined to last for 50 min similar to (Berrocal et al., 2018).

### QUANTIFICATION AND STATISTICAL ANALYSIS

#### Image Analysis of static FISH and smFISH images in Imaris

Nuclei and RNA puncta were initially detected using the Imaris software 9.2 (Bitplane, Oxford Instruments, Concord MA, RRID:SCR_007370). RNA puncta were then assigned to nuclei in a proximity based method using custom python scripts.

Nuclei were identified and segmented using the Imaris “surface” function. Nascent transcription foci were identified using Imaris “spots” function and estimated to be 0.6 µm in diameter with a z-axis point spread function of 1 µm. Single mRNA puncta were identified with spot volumes of 0.3 µm across and 0.6 µm in the z direction. False coloring of the *ush* expression domain in Figure 4Aiii was achieved using the Imaris “Cell” function. A cell mask was generated using a Spectrin antibody stain (not shown) and false colored according to the sum fluorescence of single mRNAs. Customized Python scripts were used to analyze the data extracted from Imaris and are described below.

#### Nuclear tracking and spot identification in live imaging data sets in Imaris

Nuclei were first smoothed and blurred using a wavelet filter (Imaris X-tension by Egor Zindy) and then segmented using the Imaris “surface” function based on the His-RFP fluorescent channel. Nuclei were tracked through time in 3D using the inbuilt autoregressive motion with a maximum frame gap size of 5 and a maximum travel distance of 5 µm. Active transcription sites were detected using the Imaris “spots” function in three-dimensions. Transcription foci were estimated to be 1.8 µm across with a z-axis point spread function estimation of 7.8 µm. To determine the background fluorescence of the data set, a set of “spots” was generated for background correction. Here, four spots were inserted every third time frame, avoiding nascent transcription sites. The background correction spots have the identical volume to the transcription site spots.

#### Custom python scripts for live imaging data analysis

##### 1. Spot assignment to nuclei

For both static and live imaging, spots were assigned to nuclei using the long axis of the nucleus as a reference for the midline of each nucleus. The long axis for each nucleus was calculated, using the Imaris 9.2 ellipsoid axis C and spots were then assigned to the nearest nuclei axis within the 3D space. The number of spots assigned to each nucleus was recorded.

##### 2. Nuclei distance to midline

The midline for the expression domain was calculated by fitting a polynomial (2-dimensions) using the coordinates of the mRNA spots as detected by Imaris 9.2. The distance of each nucleus was then calculated back to the midline and reported in µm.

##### 3. Mitotic Wave correction

To correct for time differences in transcriptional onset due to the mitotic wave, the temporal profile of cell areas was synchronized. The microscopy time frame at which telophase was detected was noted for each cell area along the AP axis, which was then used to set the zero time point for each position along the long axis of the embryo.

##### 4. Background subtraction

Background was recorded from the first time point where fluorescent foci were identified in the MS2 data. Background was then recorded every 3 frames until the end of the video. The background was then fit as a linear polynomial (1 dimension). The equation of the line was then used to calculate the background level at every time point. The raw value was then corrected by subtracting the background from the raw value.

All python scripts used in this analysis and instructions on use can be found at: https://github.com/TMinchington/sass

#### BMP signaling regions and 1 cell wide spatial bins

Bins of one cell width were defined to be 5 µm wide, starting at the dorsal midline. Measurements of the distance to the dorsal midline were obtained from the custom python scrips described above.

Before modeling of the transcription parameters, nuclei were grouped based on their position related to the dorsal midline. For the analysis of *ush*, the high signaling region was defined as 6 cell rows spanning the center of the expression domain (−15 to 15 µm from the dorsal midline), the medium signaling region as another three cell rows (−15 to -30 and 15 to 30 µm from the dorsal midline) and the low signaling region starting at -30 and 30 µm, respectively. To analyze *ush* in st2-*dpp* embryos, the low signaling region was defined as -30 to 50 and 30 to 50 µm with a fourth region to account for the expanded expression domain starting at -50 and 50 µm, respectively. Additionally, for the comparison of st2-*dpp* parameters to *ush* wt, only nuclei in the st2 region of *ush* wt embryos were selected for the comparison.

For the analysis of *hnt* transcription, the regions were redefined slightly due to their sparseness and narrow expression pattern. The *hnt* high signaling domain was defined as 4 cell rows spanning the dorsal midline (−10 to 10 µm) and the medium region to start at -10 and 10 µm). This resulted in the exclusion of traces from a small number of nuclei on the edges of the expression domain, which had very low transcriptional activity. Some cells were removed from the initial dataset for each embryo at several stages throughout the data processing pipeline. Cells with empty fluorescence traces and more than two missing time points were removed at the processing stage. Cells showing only low intensity, transient fluorescence activity at the edge of the expression domain were removed prior to training the model.

#### Modelling Changes in Kinetic Parameters of Transcription

We used a memory-adjusted hidden Markov model (mHMM) to infer the promoter state activity given MS2 fluorescence data (Lammers et al., 2020). The model parameters are the transition rates between on and off states of the promoter and the mean/variance of the signal in the on and off states. In order to investigate the spatial regulation of transcriptional parameters, nuclei were subdivided as described above.

The mHMM was trained separately on each of these regions per embryo in order to generate the graphs showing global transcriptional parameters per region. Inferred global transcriptional parameters included promoter switching on rate (*k*_*on*_), promoter switching off rate (*k*_*off*_), Pol II initiation rate (k_*ini*_, expressed in terms of A.U.), promoter mean occupancy (<n>), burst size (*k*_*ini*_ / *k*_*off*_) and burst frequency ((*k*_*on*_ * *k*_*off*_) / (*k*_*on*_ + *k*_*off*_)) (Zoller et al., 2018). The global parameters for each embryo were then used to generate a set of inferred posterior promoter traces for each individual cell within the embryo (using the Forward-Backward algorithm) allowing for estimation of cell-specific promoter switching rates, mean occupancy, burst frequency and amplitude.

The model state-space for the mHMM is the sequence of promoter on-off states within a window of length K which is determined by the elongation time (determined by length of the gene and estimated transcription speed, see (Lammers et al., 2020). The state-space of the mHMM is therefore 2^K^ in size. This state-space is too large for us to use the original MATLAB implementation of the model here because of computational space and time limitations, and therefore we reimplemented the model in python using a truncated state-space approximation. We used the Forward algorithm to rank states dynamically by probability given the current and previous observations in the sequence and we removed states below M in rank at each time, where M is a user-defined number of stored states that determined the accuracy of the approximation. Full details of this scalable implementation of the memory adjusted HMM are described in a forthcoming publication (Bowles et al., in preparation).

#### Single cell parameters

Single cell parameters were estimated through use of the forward-backward algorithm to infer posterior promoter state sequences for each cell within a given region of the embryo. Transition rates were calculated by counting the number of promoter state transitions for each cell, using pseudo-counts to regularise the cell-specific estimates.

#### Fluorescence to Pol II conversion

To convert the MS2 fluorescence values to Pol II molecules, the number of *ush* and *hnt* mRNAs was quantified using smFISH probes in fixed embryos of all relevant genotypes. Images were acquired from late nc14 embryos where the beginning of the cephalic furrow was visible, representing the endpoint of our live imaging data. Using the Imaris pipelines described above, the number of mRNAs per cell was quantified for cells positioned in the high BMP signaling region. We first estimated the half-lives of the *hnt* and *ush* mRNAs by using smFISH data to quantitate the number of Pol II in the transcription sites. The fluorescence of transcription sites and single mRNAs was quantified using the AirLocalize software (Trcek et al., 2017), allowing mRNA production rate to be calculated after correcting for the position of the smFISH probes (weighting times determined using the TransQuant software), and comparing this to the number of mRNAs detected in the cytoplasm (Bahar Halpern and Itzkovitz, 2016). For *ush*, the following smFISH data were used: *ush* (no MS2), *ush-MS2*, st2-*dpp* and *hnt>ush-MS2*, giving half-life values of 13.1, 10.8, 10.5 and 13.1 min, respectively. For *hnt*, we used *hnt* (no MS2), *hnt-MS2 and ush>hnt-MS2* smFISH data, from which the half-life was calculated as 3.5, 2.0 and 2.5 min, respectively. While the *ush* mRNA half-life is in the range of 6-14 min previously determined for early zygotically expressed *Drosophila* mRNAs (Boettiger and Levine, 2013; Edgar et al., 1989; Edgar et al., 1986), the *hnt* mRNA appears to be very unstable. We used half-lives of 3 and 12 min for *hnt* and *ush* mRNAs, respectively, to correct the final mRNA numbers detected by smFISH, to account for this rate of degradation. Dividing this corrected number of mRNAs produced in the imaging time window by 2 reveals the transcriptional output from a single allele. We then generated a conversion factor for the transcription traces by dividing the mean integrated fluorescent signal from nuclei in the high signaling region by the corrected mRNA output for a single allele (Lammers et al., 2020). This conversion factor was then used to scale the fluorescent signal for each embryo. We also used an alternative approach for scaling the fluorescent signals by dividing the signal averaged over three time points, equivalent to the stage in development imaged by smFISH, by the production rate estimated from Pol II number in the transcription sites (Bahar Halpern and Itzkovitz, 2016). While this approach gave similar estimates for Pol II loading rate, we find these estimates to be more variable, potentially as Pol II number in the transcription site is more difficult to accurately measure. Therefore, the fluorescent signals are converted to Pol II based on mRNA number throughout.

#### Statistical Analysis

Statistical comparisons were performed using two-tailed Student’s t tests, one-way ANOVA with multiple comparison, Pearson correlations, and paired Student’s t tests using GraphPad Prism (RRID: SCR 002798) and R (Version 3.5.2). Statistical test and sample sizes can be found in Figure legends. Statistical significance was assumed by p<0.05. Individual p values are indicated in Figure legends.

## SUPPLEMENTARY INFORMATION

**Video S1:** Maximum intensity projection of a representative embryo showing endogenous *24xMS2-ush* transcription (grey) and Histone-RFP (red) imaged with a 40x objective and 20 sec time resolution during nc14. **Related to Figure 1, S1**.

**Video S2:** As in Video S1 but showing *hnt* transcription. **Related to Figure 1, S1**.

**Video S3**: Maximum intensity projection of a representative a homozygous embryo in dorsal-lateral position with transcription from two *24xMS2-ush* alleles (grey) and Histone-RFP (red) imaged with a 40x objective at 2.15 confocal zoom and 15 sec time resolution during nc14. **Related to Figure S2. Video S4:** Maximum intensity projection of a representative embryo showing endogenous *24xMS2-ush* transcription (grey) and Histone-RFP (red) imaged with a 40x objective and 20 sec time resolution during nc14. The expression domain is broadened by a single copy of the st2-*dpp* transgene. **Related to Figure 5**.

**Video S5:** As in Video 1 but showing *ush* transcription in a *hnt>ush* embryo. **Related to Figure 6, S6**.

**Video S6:** As in Video 1 but showing *hnt* transcription in a *ush>hnt* embryo. **Related to Figure 7, S7**.

**Table 1: smFISH probes used for FISH and complementary to exonic sequences in *hnt* and**

***ush*, related to STAR Methods**.

---

**Hoppe_Figure S1.**
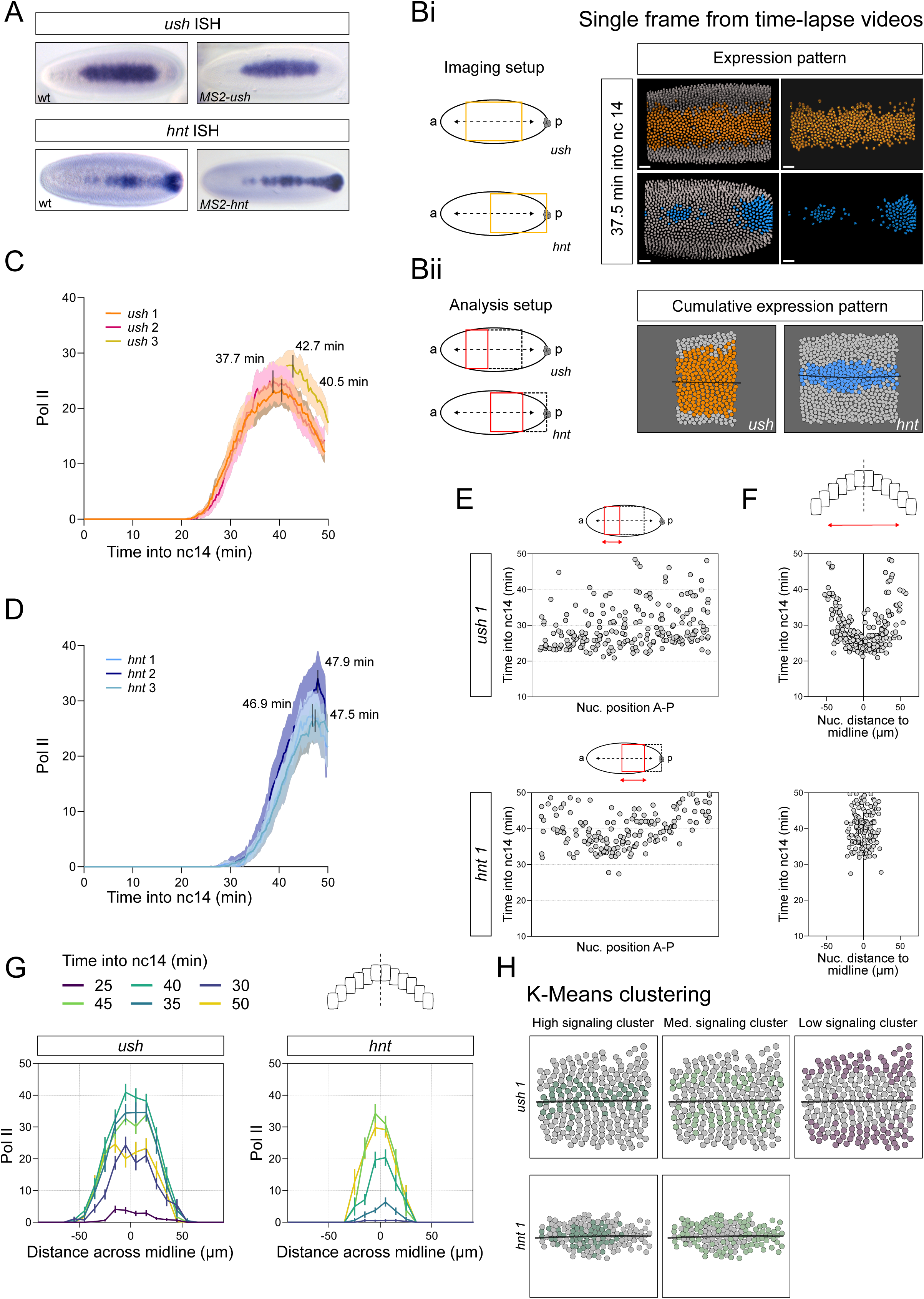

**Hoppe_Figure S2.**
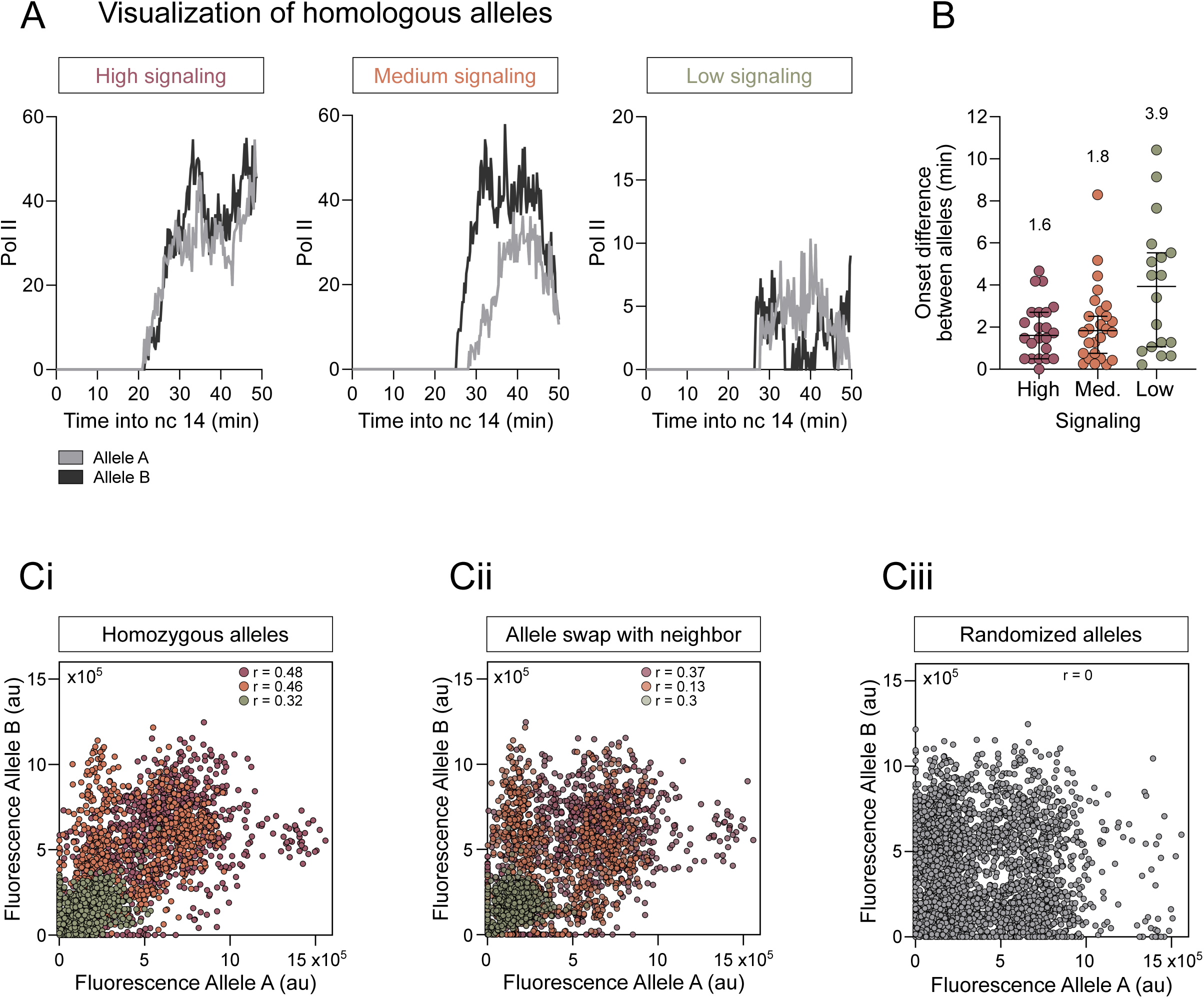

**Hoppe_Figure S3.**
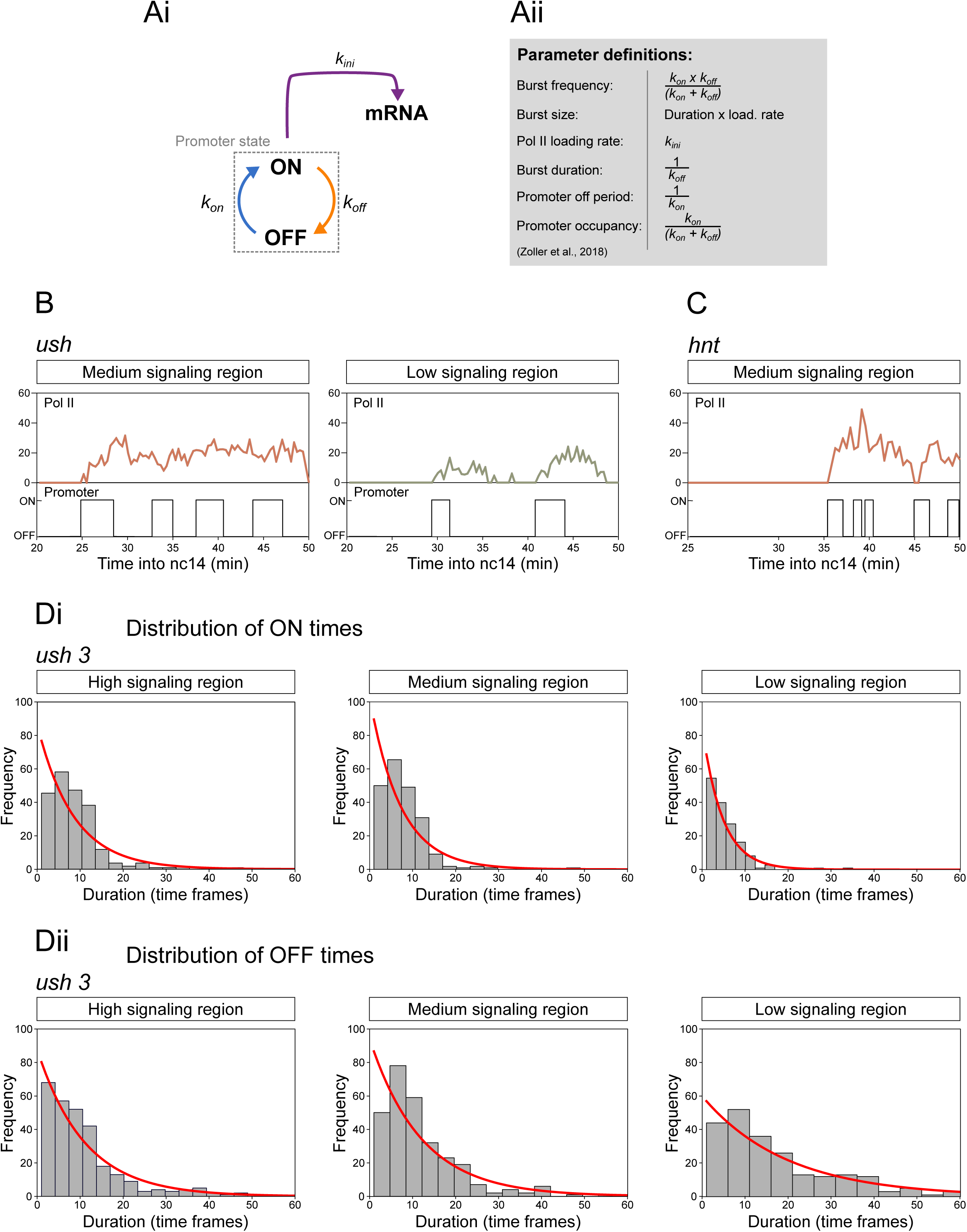

**Hoppe_Figure S4.**
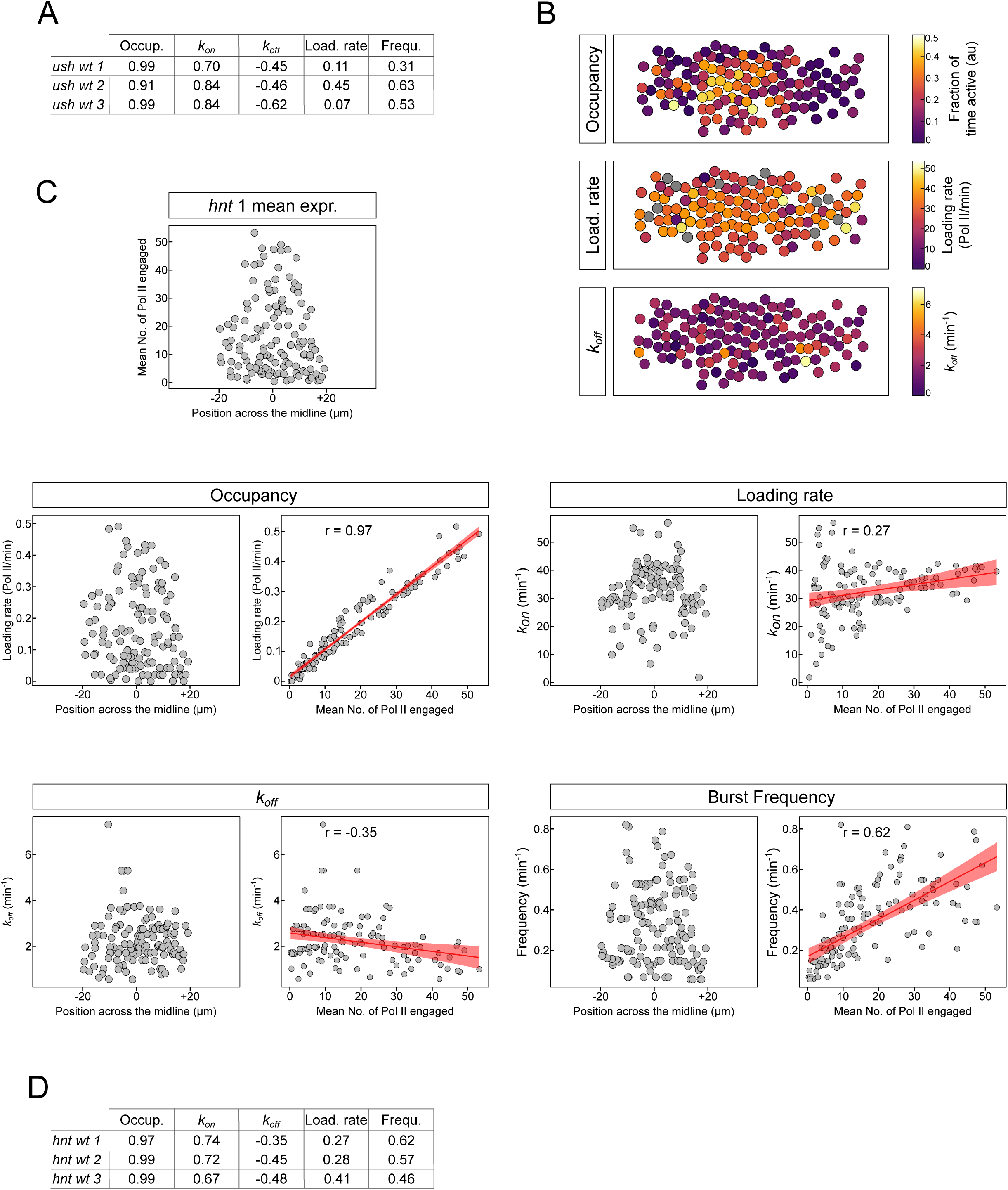

**Hoppe_Figure S5.**
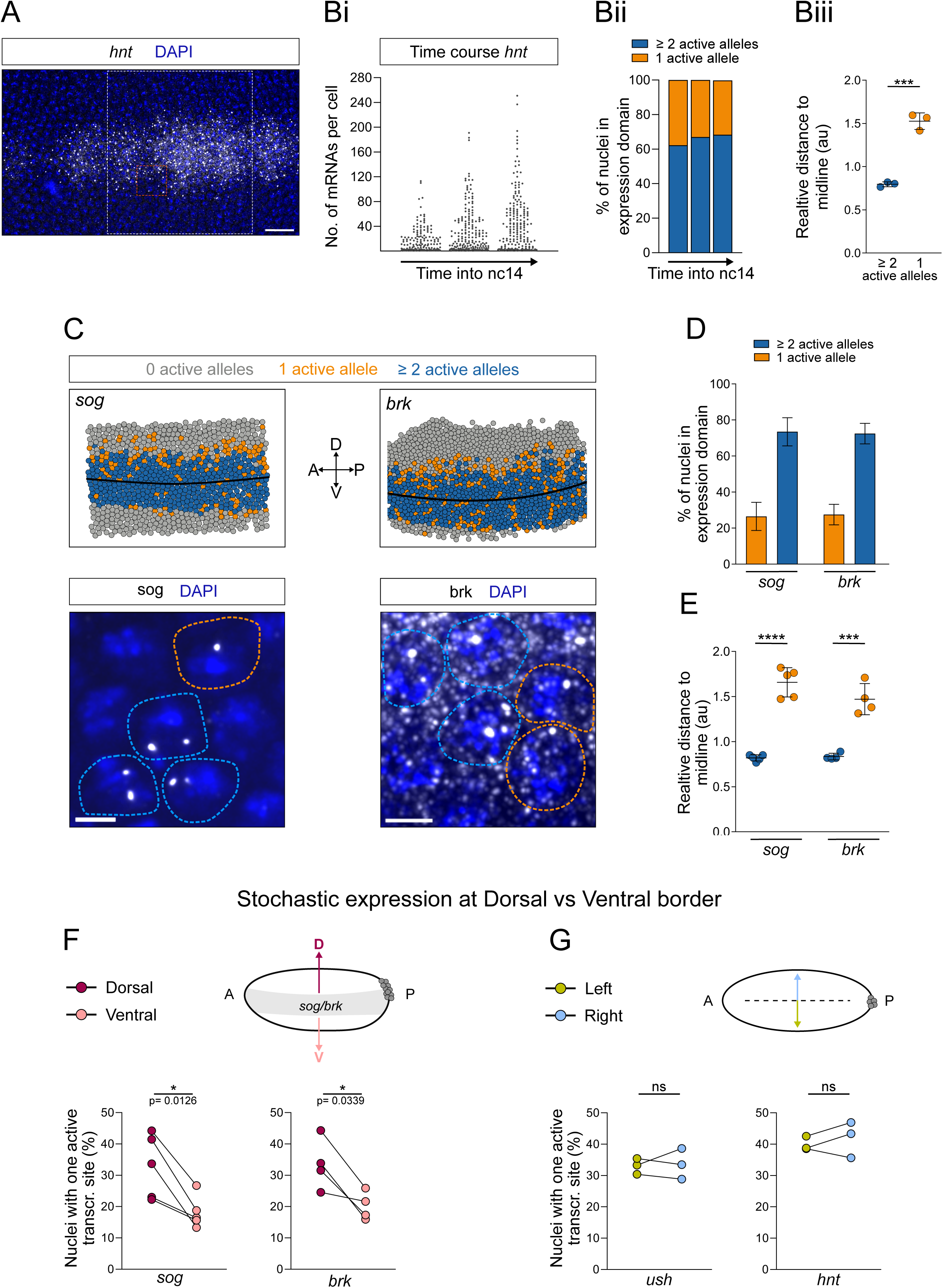

**Hoppe_Figure S6.**
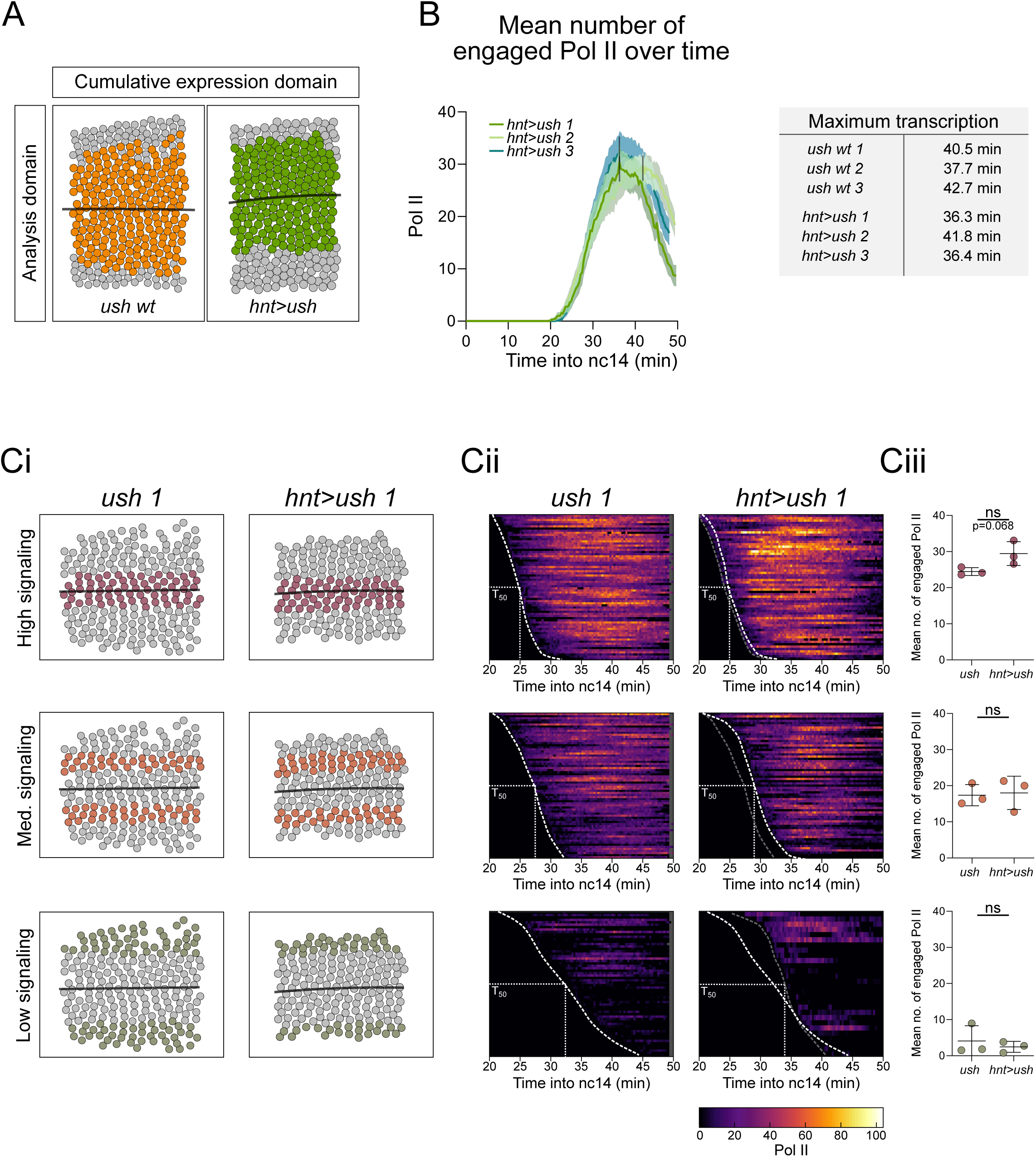

**Hoppe_Figure S7.**
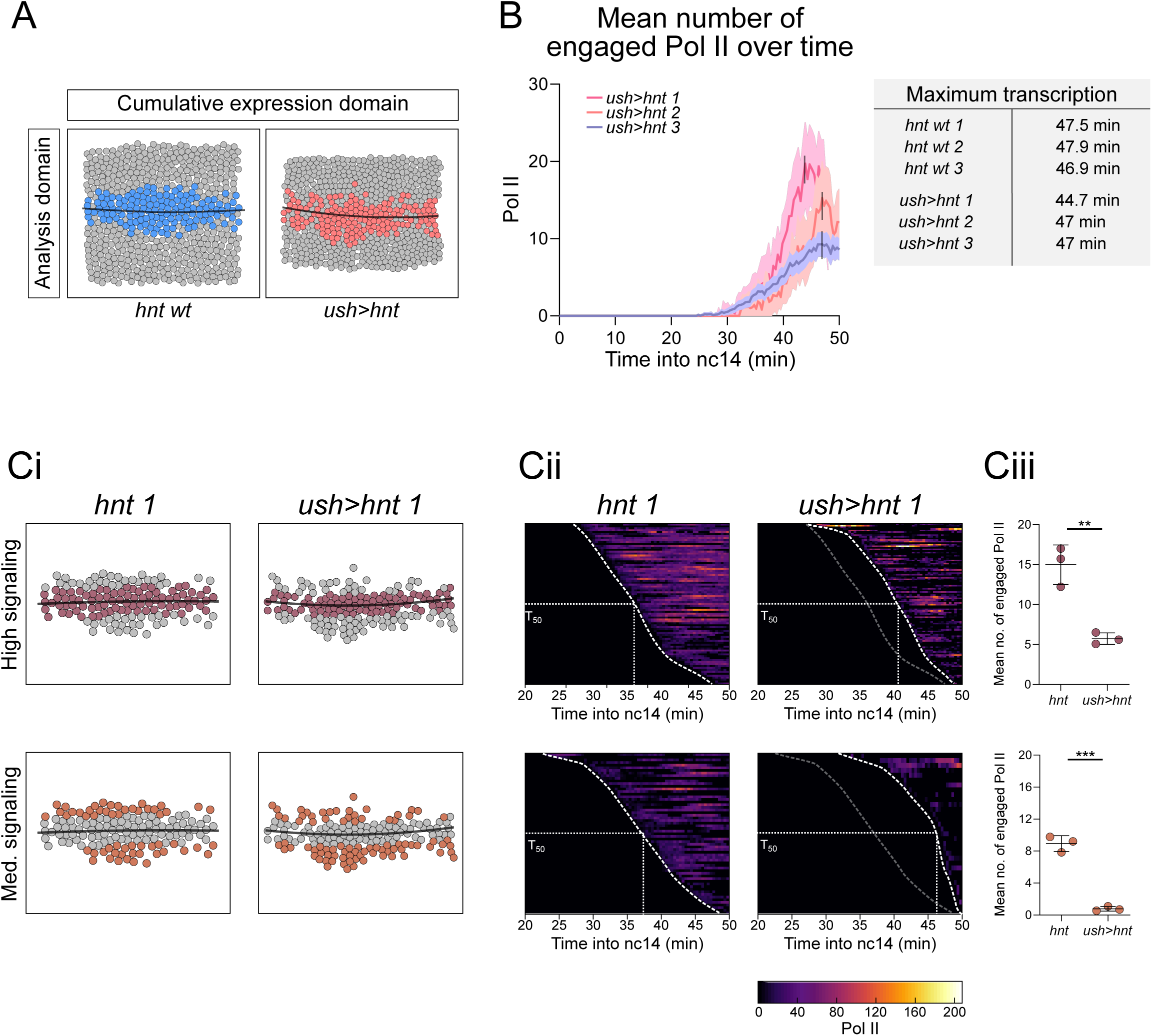

